# Hybrid offspring of C57BL/6J mice exhibit improved properties for neurobehavioral research

**DOI:** 10.1101/2022.05.03.490527

**Authors:** Hadas E. Sloin, Lior Bikovski, Amir Levi, Ortal Amber-Vitos, Tomer Katz, Lidor Spivak, Shirly Someck, Roni Gattegno, Shir Sivroni, Lucas Sjulson, Eran Stark

## Abstract

C57BL/6 is the most commonly used mouse strain in neurobehavioral research, serving as a background for multiple transgenic lines. However, C57BL/6 exhibit behavioral and sensorimotor disadvantages that worsen with age. We bred FVB/NJ females and C57BL/6J males to generate first-generation hybrid offspring, (FVB/NJ x C57BL/6J)F1. The hybrid mice exhibit reduced anxiety-like behavior, improved learning, and enhanced long-term spatial memory. In contrast to both progenitors, older hybrids maintain sensorimotor performance and exhibit improved long-term memory. The hybrids are larger than C57BL/6J, exhibiting enhanced running behavior on a linear track during freely-moving electrophysiological recordings. Hybrids exhibit typical rate and phase coding of space by CA1 pyramidal cells. Hybrids generated by crossing FVB/NJ females with transgenic males of a C57BL/6 background support optogenetic neuronal control in neocortex and hippocampus. The hybrid mice provide an improved model for neurobehavioral studies combining complex behavior, electrophysiology, and genetic tools readily available in C57BL/6 mice.

## Introduction

Many advances in behavioral and biomedical research rely upon lab animals. In choosing an animal model, there are always two conflicting considerations. On the one hand, an organism as similar as possible to the object of interest is desired. On the other hand, the least advanced organism for answering the research question is preferred. The lab mouse (*Mus musculus*) has emerged as a key model in balancing these demands, sharing multiple physiological systems and genes with humans while exhibiting simple mating. Over the years, numerous inbred mouse strains have been developed, allowing genetic modifications, labeling, and manipulation of specific proteins, cells, and organs (Beck et al., 2000). However, there are phenotypic differences between inbred strains used as background for transgenic mice (Upchurch and Wehner, 1988; Võikar et al., 2001; Wahlsten et al., 2003; Brown and Wong, 2007; O’Leary et al., 2011; O’Leary et al., 2013; Kafkafi et al., 2017). Hence, mouse strain selection is essential when designing a scientific project.

The mouse strain used most often in neurobehavioral research is C57BL/6. The C57BL/6 Jackson Laboratory substrain, C57BL/6J, provided the first extensively sequenced mouse genome (Waterston et al., 2002) and is among the most widely used inbred strains (Altman and Katz, 1979; Mekada et al., 2009). An advantage of inbreeding is that differences between individuals of the same strain are minimal. The genetic similarity of inbred animals is especially useful for knockout studies, which may require homozygous animals (Silva et al., 1997). Indeed, a large variety of transgenic mice on the C57BL/6 background is available. However, inbreeding exposes undesired phenotypical traits due to homozygous recessive alleles, and does not guarantee stability over generations due to genetic drift (Brekke et al., 2018). C57BL/6-derived mice exhibit known phenotypic disadvantages, including sensitivity to pain, addiction, impaired balance, age-dependent hearing loss, and increased anxiety (Mogil et al., 1999; Crawley, 1996; Ouagazzal et al., 2006). Thus, the behavioral tasks studied using C57BL/6 are limited.

One way to balance the requirements of convenient genetic control and complex behavior is to generate hybrids of C57BL/6J and another strain. Hybrids inherit one allele from the C57BL/6J parent, maintaining transgenic properties in a heterozygous manner. Previous work has shown that offspring of C57BL/6J (C57) and FVB/NJ (FVB) mice consume more alcohol than either progenitor, serving as a preferable model for alcohol consumption (Blednov et al., 2005, 2010). FVB is an inbred albino strain which carries a recessive allele causing retinal degeneration (Pittler and Baehr, 1991; Taketo et al., 1991). The C57 and FVB strains have distinct genealogies (Beck et al., 2000; Mekada et al., 2009), increasing genetic heterogeneity of the hybrid offspring. Thus, hybrids may serve as a potentially favorable animal model.

Here, we bred C57 males with FVB females to determine whether the first-generation hybrids (HYB) are a preferable model for neurobehavioral studies. We tested mice of the three strains (C57, FVB, and HYB) using a battery of standard phenotyping assays (Crawley, 2008; Fuchs et al., 2011). Compared to either inbred strain, HYB exhibited similar sensorimotor performance, reduced anxiety-like behavior, faster learning, and improved memory, which were maintained upon aging. HYB mice were physically larger, and during electrophysiological recordings exhibited enhanced running on a linear track compared to C57, while CA1 neurons had similar place coding properties. Furthermore, transgenic HYB supported optogenetic neuronal control in neocortex and CA1. Together, the results suggest that the hybrid mice are a preferable model for systems neuroscience studies combining behavioral tests, electrophysiology, and genetic targeting.

## Results

### Hybrid mice are larger than C57 mice and exhibit similar sensorimotor performance

We decided to use offspring of C57 and FVB because the two inbred strains are of different genealogies (Beck et al., 2000) and exhibit distinct phenotypic disadvantages (Crawley, 2008). FVB dams produce large litters (Silver, 1995) and were therefore selected as the maternal strain. To maximize the genetic similarity between individual mice, we focused on (FVB/NJ x C57BL/6J)F1, the first-generation offspring (HYB; **Table S1**). We found that sedentary HYB were larger than either parental strain. At the age of 9 months, the median [interquartile interval, IQR] weight of HYB was 45.4 [41.8 47.2] g, compared to 30.3 [29.3 32.2] g for C57 (*p*=9.6×10^-10^; Kruskal-Wallis test, corrected for multiple comparisons; **Fig. 1**, **Fig S1A**, and **Table S2**). Higher weights are beneficial for electrophysiological experiments in freely-moving animals, in which animal size limits the weight of the implanted apparatus. Examination of spontaneous home cage behavior (**Fig. S1B**) and gait analysis (**Fig. S1C**) revealed consistent differences between the three strains. Thus, the HYB constitute a unique, larger strain.

**Figure 1:**
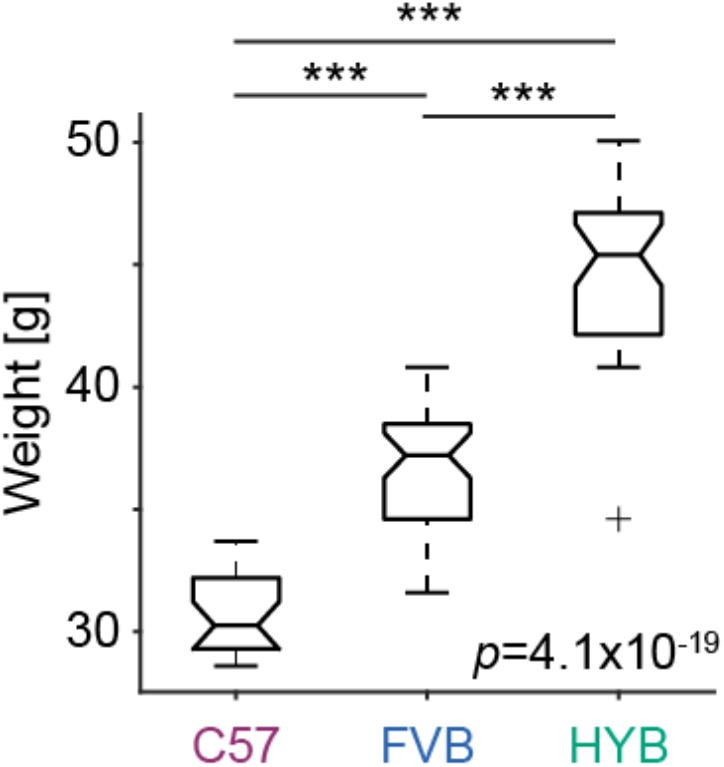
Hybrid mice weigh more than both progenitors. Mice of the three strains were weighted periodically at ages between 255 and 285 days (approximately 9-month-old), and weights were averaged over measurements for every mouse. Every box plot shows median and interquartile range (IQR), whiskers extend for 1.5 times the IQR in every direction, and a plus indicates an outlier. *P*-value indicated by text is for a one-way Kruskal-Wallis test. ***: *p*<0.001, Kruskal-Wallis test, corrected for multiple comparisons.

To assess the sensorimotor capabilities of HYB, we subjected 9-month-old mice to three standard assays (n=19 C57, 14 FVB, and 19 HYB; **Table S1** and **Table S2**). Visual behavior was quantified in an optomotor drum test (Abdeljalil et al., 2005), in which mice follow vertical visual stimuli by making directional head movements. C57 and HYB made similar numbers of head turns (C57: 8 [5 11] HYB: 8 [5 9], *p*=0.96, Kruskal-Wallis test; **Fig. S2A**). In contrast, FVB did not make any head turns (0 [0 0]; C57 vs. FVB: *p*=1.2×10^-6^; HYB vs. FVB: *p*=4.7×10^-6^), consistent with blindness due to retinal degeneration (Pittler and Baehr, 1991). Motor endurance was tested by running the mice on a treadmill at gradually increasing speed, and measuring the distance to failure (Marques-Aleixo et al., 2015). HYB stayed on the treadmill for 126 [59 164] s, not consistently different from C57 (91 [49 120] s; *p*=0.07; Kruskal-Wallis test), but longer than FVB (40.5 [29 64] s; *p*=0.002; **Fig. S2B**). Balance was assessed by placing the mice on an accelerating rotarod for five consecutive days, and measuring the latency to fall (Jones and Roberts, 1968). Latency to fall increased over days for all strains, indicative of balance learning (C57: *p*=4.3×10^-10^; FVB: *p*=0.008; HYB: *p*=3.9×10^-10^; intra-strain Kruskal-Wallis tests; **Fig. S2C**). Following learning, C57 and HYB exhibited longer latency to fall compared to FVB (C57: 208 [197 230] s; FVB: 143 [100 196] s; HYB: 246 [183 277] s; *p*=0.03 and *p*=0.0004, Kruskal-Wallis test). To summarize, at the age of 9 months, sensorimotor performance of HYB is improved compared to FVB and similar to that of C57.

### Hybrid mice exhibit reduced anxiety and improved learning and memory

To characterize neuropsychiatric phenotypes and cognitive performance, we tested 9-month-old mice using five assays (**Table S1**). Anxiety-like behavior was studied using the elevated plus maze (Rodgers and Dalvi, 1997) and quantified by a “contra-anxiety index” that ranges from −1 to 1, defined as time spent in (open-closed)/(open+closed) arms. Indices closer to one indicate that more time was spent in the two “open” arms, corresponding to reduced anxiety-like behavior. C57 spent consistently more time in the closed arms compared to the open arms (−0.56 [−0.75 −0.46]; *p*=0.00013, Wilcoxon’s signed-rank test), while HYB did not (−0.15 [-0.27 0.15], *p*=0.21; **Fig. 2A** and **Fig. S3A**). Correspondingly, HYB spent more time in the open arms than C57, indicating reduced anxiety-like behavior (*p*=4.1×10^-6^; Kruskal-Wallis test; **Fig. 2A** and **Table S2**).

**Figure 2:**
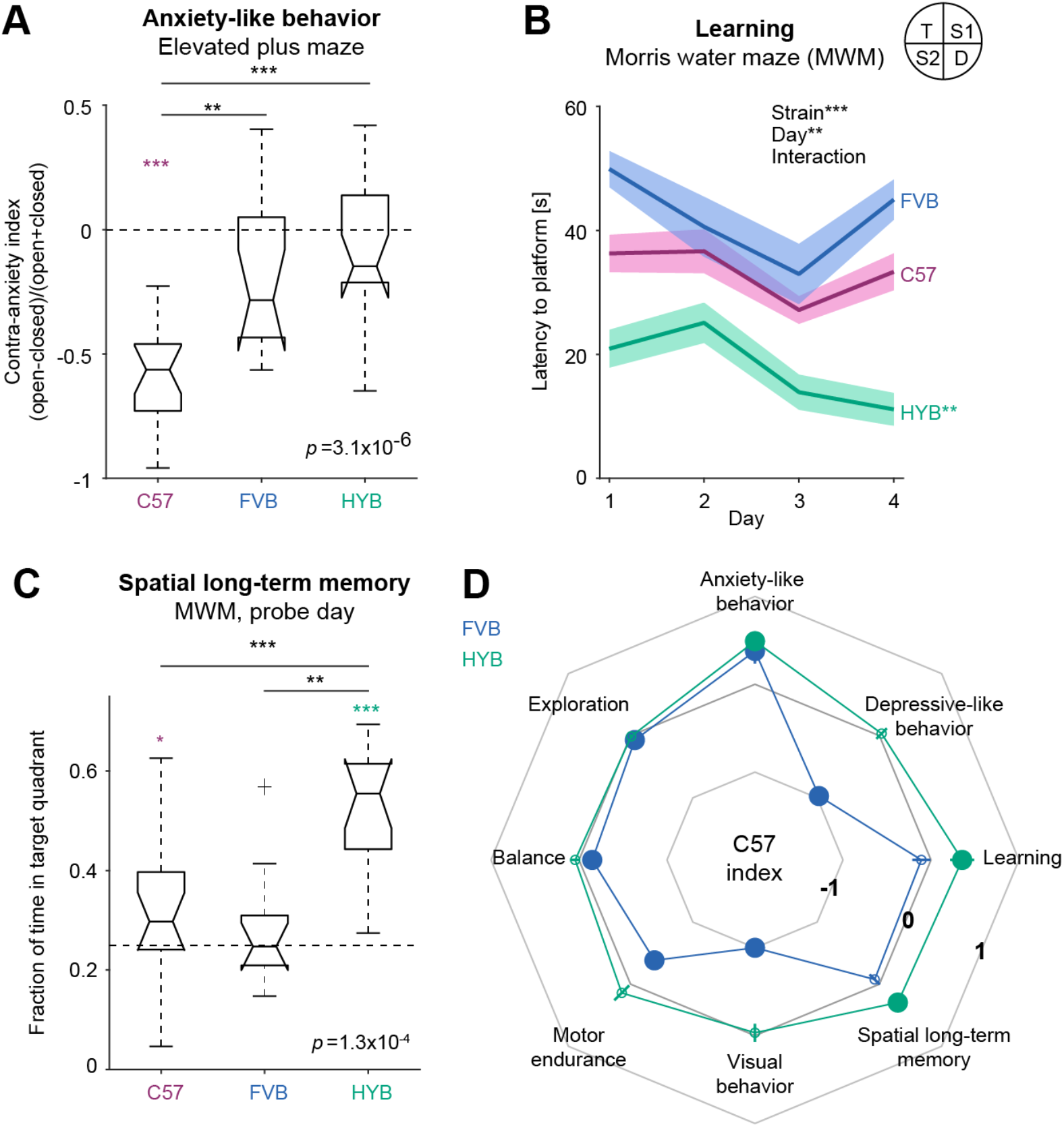
Hybrid mice exhibit reduced anxiety and improved learning and memory. **(A)** At the age of 9 months, HYB mice exhibit reduced anxiety-like behavior compared to C57. Anxiety-like behavior was tested in an elevated plus maze, and quantified using a “contra-anxiety index”, defined as the time spent in the open minus the time spent in the closed arms, divided by the sum. Lined **/***: *p*<0.01/*p*<0.001, Kruskal-Wallis test. Here and in **C**, */***: *p*<0.05/*p*<0.001, Wilcoxon’s signed-rank test, comparing to chance level (horizontal dashed line). **(B)** HYB exhibit improved learning in the Morris water maze (MWM) task compared to C57 and FVB, indicated by decreasing latency to the platform over days. The platform was submerged in the target quadrant (T). Bands show mean and SEM; **/*** next to the text indicate *p*<0.01/*p*<0.001 for a two-way, strain by day, Kruskal-Wallis test; ** at right indicates *p*<0.01, for a within-strain, across-day oneway Kruskal-Wallis test. **(C)** Compared to C57 and FVB, 9-month-old HYB exhibit improved spatial longterm memory, indicated by longer fractions of time spent in the target quadrant during the probe day. All conventions are the same as in **A**. **(D)** Comparison of HYB (or FVB) to C57 performance in eight behavioral assays using the “C57 index” at the age of 9 months. Positive Indices indicate improved performance compared to C57. Circles (and lines) at vertices show mean (and SEM) indices; filled circles indicate significant changes compared to C57 (*p*<0.05, bootstrap test).

Depressive-like behavior was tested using a forced swim test, in which longer freeze duration indicates behavioral despair (Porsolt et al., 1977). HYB and C57 exhibited similar freeze durations (C57: 130 [86 172] s; HYB: 138 [85 189] s; *p*=0.95, Kruskal-Wallis test; **Fig. S3B** and **Table S2**). Motivational behavior was quantified by activity levels in an open field (Christmas and Maxwell, 1970). We did not observe any consistent differences between C57 and HYB in overall activity in the field (C57: 0.35 [0.33 0.38]%; HYB: 0.35 [0.28 0.37]%; *p*=0.66, Kruskal-Wallis test; **Fig. S3C** and **Table S2**). Thus, compared to C57, HYB exhibit reduced anxiety-like behavior, similar depressive-like behavior, and similar motivational behavior.

To assess learning and memory behavior, we trained 9-month-old mice on the Morris water maze (MWM) task (Morris, 1981; n=18 C57, 11 FVB, and 15 HYB; **Table S1**). During four consecutive days, mice were placed in a pool with an invisible submerged platform. Over days, only HYB exhibited a decrease in the latency to platform, indicative of learning (C57: *p*=0.096, FVB: *p*=0.053, HYB: *p*=0.008, intrastrain Kruskal-Wallis tests; **Fig. 2B**). Pooled over days, the mean±SEM latency to the platform was shortest for the HYB (17.8±3 s), compared to C57 (33.3±2.9 s) and FVB (42.1±4 s; *p*=1.7×10^-13^, Kruskal-Wallis test; **Table S2**). Similar results were obtained when examining the distance traveled (Terry, 2009) instead of the latency to platform (**Fig. S3D**). Hence, compared to both progenitors, HYB exhibit improved learning performance.

To assess spatial long-term memory, the mice were tested in the MWM during a “probe” day, without a platform. HYB spent over half of the time in the target quadrant, indicative of spatial long-term memory (median [IQR] fraction of time: 0.55 [0.36 0.62]; *p*=6.1×10^-5^, Wilcoxon’s signed-rank test compared to chance level, 0.25; **Fig. 2C**, **Fig. S3Ea**, and **Table S2**). Accordingly, HYB spent a larger fraction of time in the target quadrant than C57 (0.3 [0.24 0.4], *p*=0.002; Kruskal-Wallis test) and FVB (0.25 [0.15 0.31], *p*=0.0003; **Fig. 2C** and **Fig. S3Eb**). Similar results were obtained when spatial long-term memory was quantified by the fraction of visits to the target quadrant (*p*=0.0014, Kruskal-Wallis test; **Fig. S3Ec**) or by the latency to the platform (*p=*3.7×10^-3^, Kruskal-Wallis test; **Fig. S3Ed**). Thus, in contrast to both progenitors, HYB exhibit robust spatial long-term memory behavior.

To quantify differences between HYB (or FVB) and the C57, we treated the C57 strain as a reference and computed “C57 indices” for the eight assays (**Fig. 2D**). For a given metric, the C57 index is defined as the difference between the median value of HYB (or FVB) and the median value of C57, divided by the sum. Therefore, a C57 index above zero indicates higher values of the metric for the tested strain, compared to C57 performance. HYB performance was similar to C57 in assays testing visual behavior (mean±SEM C57 index: −0.031±0.1, *p*=0.34, bootstrap test), motor endurance (0.14±0.12, *p*=0.09), balance (0.04±0.05, *p*=0.14), exploration (−0.01±0.023, *p*=0.35), and depressive-like behavior (0.04±0.08, *p*=0.32). In contrast, HYB exhibited reduced anxiety-like behavior (0.49±0.1, *p*=0.00001), improved learning (0.35±0.14, *p*=0.002), and improved spatial long-term memory (0.3±0.06, *p*=0.0003). Thus, 9-month-old HYB exhibit sensorimotor properties similar to C57, while exhibiting reduced anxiety-like behavior and improved learning and memory performance.

### Older hybrids maintain sensorimotor performance and exhibit improved memory

To examine the suitability of the HYB for long-term studies that require stable behavioral performance, we applied all assays also to 3-month-old mice (**Table S1**). To quantify performance differences between 3-month-old and 9-month-old mice of the same strain, we defined an “aging index” (**Fig. 3**). For a given performance metric, the aging index is the difference between the median value obtained for 9-month-old and 3-month-old mice, divided by the sum. Thus, an aging index of zero indicates no age-related changes in performance, whereas positive indices indicate age-dependent increases. Visual behavior was maintained across age groups for all strains (mean±SEM aging indices: C57, 0.03±0.17, *p*=0.41; FVB, not available; HYB, 0.04±0.13, *p*=0.28, bootstrap test; **Fig. 3Aa**). While C57 and FVB exhibited reduced motor endurance upon aging (C57: −0.41±0.1, *p*=0.0001; FVB: −0.5±0.13, *p*=0.0014), HYB endurance was maintained (−0.23±0.22, *p*=0.11; **Fig. 3Ab**). Finally, while C57 and FVB balance performance declined between age groups (C57: −0.2±0.06, *p*<0.05; FVB: −0.31±0.09, *p*<0.05), HYB balance was maintained (0.1±0.13, *p*=0.75; **Fig. 3Ac**). Thus, while the motor capabilities of C57 and FVB deteriorate upon aging, HYB motor performance remains stable across the two age groups studied.

**Figure 3:**
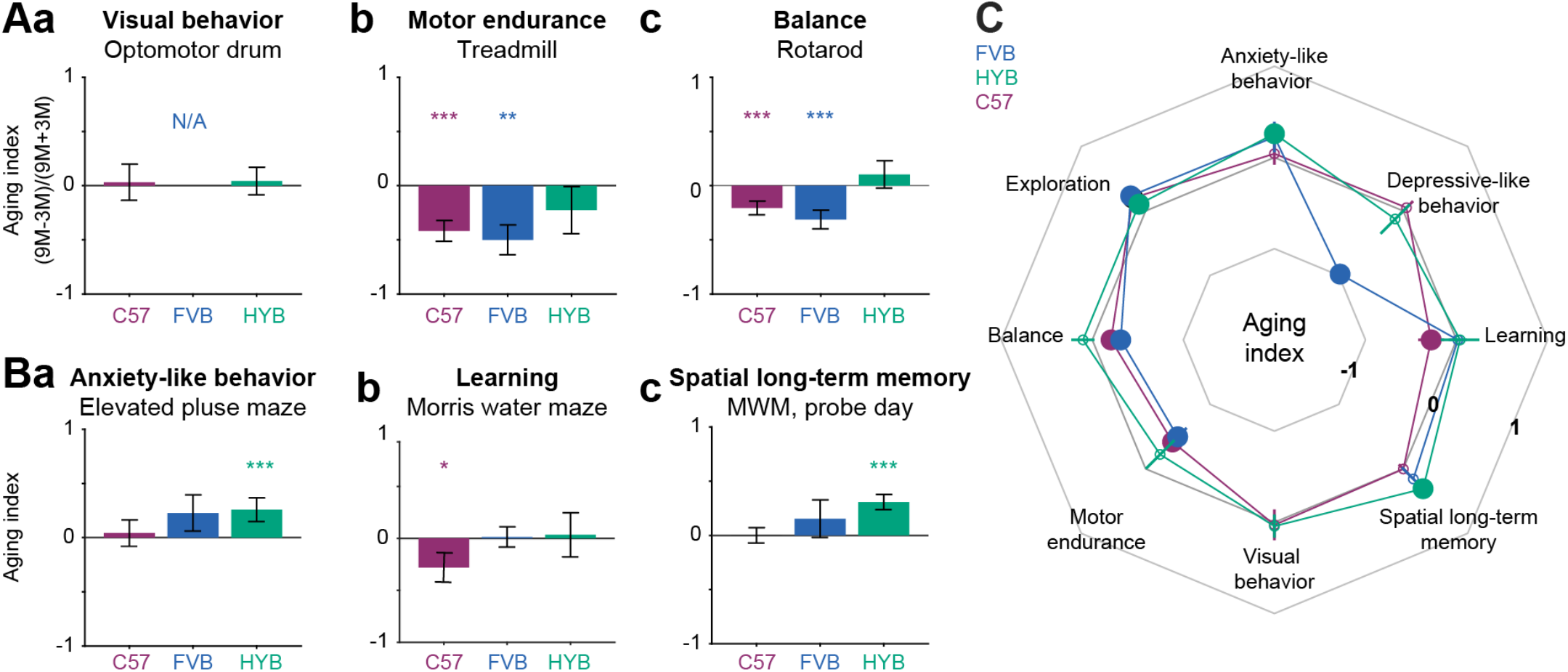
Older hybrids maintain sensorimotor performance and exhibit improved memory. **(A)** Sensorimotor behavior remains intact at an older age for HYB but not for C57 or FVB mice. Positive aging indices indicate improvement across 3-month-old and 9-month-old mice of the same strain. (**a**) Visual behavior of HYB and C57 remains intact at an older age, whereas FVB do not react to rotating visual stimuli at any age. Here and in **B**, bars indicate mean; error bars, SEM; */**/***: *p*<0.05/*p*<0.01/*p*<0.001, bootstrap test compared to a no-change null. (**b**) Motor endurance is reduced for older C57 and FVB and maintained for older HYB. (**c**) Balance performance deteriorates for C57 and FVB upon aging and is maintained for older HYB. **(B)** Psychiatric-like and cognitive performance across age groups. All conventions are the same as in **A**. (**a**) HYB, but not C57 or FVB, exhibit reduced anxiety-like behavior upon aging. (**b**) Learning performance of C57, but not FVB or HYB, deteriorates upon aging. (**c**) Spatial longterm memory performance of C57 and FVB is maintained upon aging, whereas HYB memory performance improves. **(C)** Comparison of performance changes in eight behavioral assays using the “Aging index”. All conventions are the same as in **Fig. 2D**.

At the age of 9 months, HYB exhibited reduced anxiety-like behavior and improved learning and memory performance compared to C57 (**Fig. 2D**). We tested whether performance on the three assays was maintained across age-groups in every strain. Compared to 3-month-old mice of the same strain, anxiety-like behavior was reduced for 9-month-old HYB (mean±SEM aging index: 0.26±0.11, *p*=0.0005, bootstrap test), but not for C57 (0.04±0.12, *p*=0.37) or FVB (0.22±0.17, *p*=0.07; **Fig. 3Ba**). Learning performance was reduced in older C57 mice (−0.28±0.14, *p*=0.03), but maintained for older HYB (0.04±0.21, *p*=0.43; **Fig. 3Bb**). Both C57 and FVB maintained spatial long-term memory performance between age groups (C57: 0.008±0.13, *p*=0.28; FVB: −0.07±0.16, *p*=0.16). In contrast, older HYB exhibited improved spatial long-term memory (0.31±0.07; *p*=0.0003; **Fig. 3Bc**). Thus, anxiety-like behavior is reduced and memory performance improves upon aging in HYB (**Fig. 3C**).

### Hybrid mice exhibit enhanced linear track running and typical CA1 place coding

The results reported in **Figs. 1–3** show that HYB exhibit certain advantages over C57. However, improved performance on behavioral assays spanning several minutes does not necessarily predict enhanced behavior over daily recording sessions that span hours and include the additional weight of an implanted apparatus. To assess the suitability of freely-moving HYB for electrophysiological experiments, we performed high-density recordings from two HYB and two C57 mice running back and forth on a 150 cm linear track (**Table S3**). Behavior in the first 14 sessions on the track was assessed to quantify performance at similar experience levels. HYB carried out a median [IQR] of 164 [139 215] one-direction trials per session, while C57 ran 136 [113 178] trials (*p*=0.015, Mann-Whitney’s *U*-test; **Fig. 4A** and **Table S4**). Furthermore, HYB exhibited higher running speeds, 31 [22 44] cm/s compared to 23 [18 30] cm/s for C57 (*p*=0.007, *U*-test). Thus, freely-moving HYB exhibit enhanced motor abilities and exploration during prolonged electrophysiological recordings.

**Figure 4:**
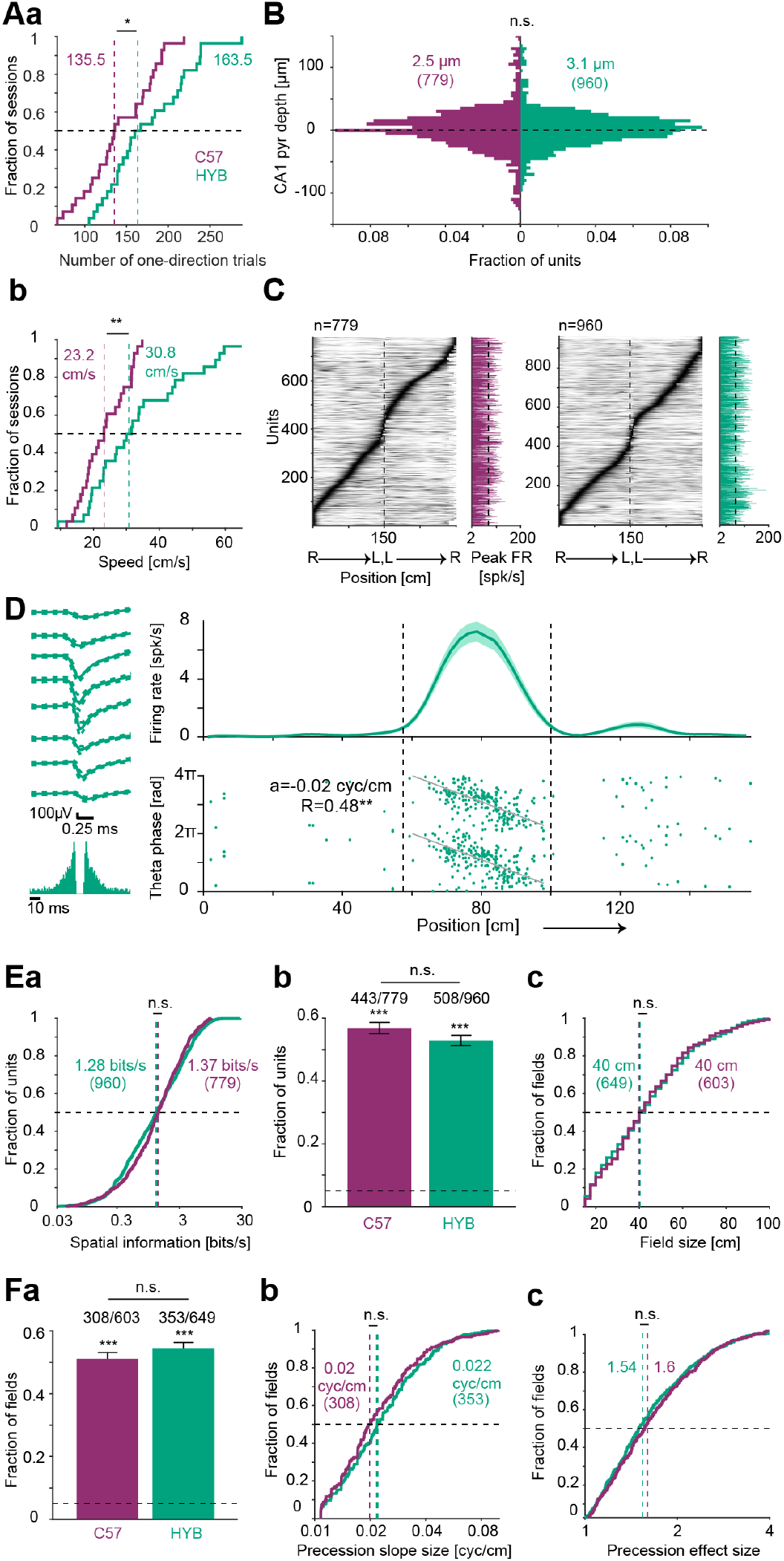
Hybrid mice exhibit enhanced linear track running and typical CA1 place coding. **(A)** Compared to C57 mice, HYB carry out more trials and run faster on a 150 cm long linear track. Only the first 14 sessions from every mouse were employed (two C57 and two HYB mice). */**: *p*<0.05/*p*<0.01, *U*-test. Dashed vertical lines indicate group medians. (**a**) The number of one-direction trials. (**b**) Mean running speed over trials. **(B)** Location of PYR somata relative to the center of the CA1 pyramidal cell layer (dashed horizontal line) is not consistently different between strains. The estimated depths of PYR recorded by high-density silicon probes were partitioned into 5 μm bins. Positive numbers correspond PYR closer to str. oriens. n.s.: *p*>0.05, *U*-test. **(C)** Units recorded from both strains exhibit increased firing rates within specific regions of the linear track. Each row represents a unit; firing rates on right (R) to left (L) runs are concatenated with L to R runs and scaled to the 0-1 (white-black) range for presentation purposes. Bar graphs (*right*) show peak on-track firing rates. **(D)** PYR recorded from a HYB exhibits typical place field and theta phase precession. *Top*: Firing rate as a function of position (mean±SEM over 82 right-to-left trials). *Bottom*: Theta phase and animal position at the time of every spike; phase of zero corresponds to theta peak. Running direction is presented from left to right, and vertical dashed lines indicate place field limits. *a* represents phase precession slope, and *R* represents the goodness of fit of spikes to the precession model; **: *p*<0.01, permutation test. *Left*: Wide-band (0.1-7,500 Hz) spike waveforms (mean±SD) recorded on eight consecutive sites, vertically spaced by 20 μm. *Bottom left*: Autocorrelation histogram. **(E)** HYB and C57 exhibit similar CA1 spatial rate coding. (**a**) Spatial information rate. n.s.: *p*>0.05, *U*-test. Here and in **c**, vertical dashed lines indicate group medians. (**b**) Fraction of CA1 PYR with one or more place fields out of all units active and stable on the track; n.s.: *p*>0.05, *G*-test. Here and in **Fa**, ***: *p*<0.001, binomial test, compared to chance level (0.05; horizontal dashed line). Error bars, SEM. **(c)** Place field size; n.s.: *p*>0.05, *U*-test. **(F)** HYB and C57 exhibit similar CA1 spatial phase coding. (**a**) Fraction of fields exhibiting theta phase precession; n.s.: *p*>0.05, *G*-test. (**b**) Precession slope size. (**c**) Precession effect size; n.s.: *p*>0.05, *U*-test.

To examine whether CAl pyramidal neurons (PYR) exhibit similar place coding in HYB and C57 mice tested under the same conditions, we analyzed the activity of CAl pyramidal layer PYR active on the track (C57: n=779, HYB: n=960; **Fig. 4B** and **Table S4**). In both strains, most active PYR exhibited increased firing rates in specific parts of the track (**Fig. 4C**). An example PYR recorded in HYB exhibited typical spatial rate coding (place field; O’Keefe and Dostrovsky, 1971; Moser et al., 2008) and spatial phase coding (theta phase precession; O’Keefe and Recce, 1993; **Fig. 4D**). At the group level, there was no consistent difference in the spatial information carried by PYR, with 1.28 [0.47 3.18] bits/s for HYB PYR and 1.37 [0.63 2.87] bits/s for C57 PYR (*p*=0.56, *U*-test; **Fig. 4Ea**). Place fields were detected in 443/779 (57%) of C57 PYR and in 508/960 (53%) of HYB PYR (**Fig. 4Eb**). Both fractions were above expected by chance (*p*<1.11×10^-16^, exact binomial test) and were not consistently different from each other (*p*=0.37, *G*-test). Furthermore, place field sizes were not consistently different, spanning 40 [27.5 57.5] cm for C57 PYR and 40 [25 57.5] cm for HYB PYR (603 C57 fields and 649 HYB fields; *p*=1, *U*-test; **Fig. 4Ec**). Thus, CA1 PYR recorded in HYB and in C57 mice exhibit similar rate coding of space.

To determine whether HYB CA1 PYR exhibit typical phase coding of space, we examined theta phase precession within PYR place fields. Of all fields, 308/603 (51%) C57 PYR and 353/649 (54%) HYB PYR exhibited consistent spatial theta phase precession (*p*<1.11×10^-16^, binomial test; **Fig. 4Fa**). The fraction of place fields with precession was not consistently different between the two strains (*p*=0.51, *G*-test). Precession slope size was not consistently different between strains, being 0.02 [0.01 0.03] cyc/cm for C57 PYR place fields and 0.02 [0.02 0.03] cyc/cm for HYB PYR place fields (*p*=0.071, *U*-test; **Fig. 4Fb**). Finally, precession effect size was not consistently different between strains, with 1.6 [1.28 2.13] for C57 phase precessing fields and 1.54 [1.24 2.06] for HYB fields (*p*=0.29, *U*-test; **Fig. 4Fc**). Thus, CA1 PYR recorded in HYB and in C57 exhibit similar phase coding of space.

### Transgenic hybrid mice enable optogenetic control of individual neurons

A major advantage of C57 mice for neurobehavioral research is the myriad transgenic strains available on the same background. To combine genetic control available for C57 with the enhanced behavior of the HYB, we first generated dual-transgenic mice on a C57 background by breeding CaMKII-Cre females with Ai32 males. We then crossed the male offspring with FVB females. The resulting dual-transgenic HYB are susceptible to blue-light optogenetic activation of projection neurons, including neocortical and CA1 PYR. Optogenetic PYR activation was carried out in transgenic HYB implanted with multi-site optoelectronic probes (**Table S3**). A total of 5,568 PYR and 1,066 putative interneurons (INT) were recorded from the CA1 pyramidal layer during n=55 sessions (**Table S5**). In every recording session, we tested the response of the recorded units to 50 ms blue-light pulses. During illumination, PYR exhibited increased firing rates (**Fig. 5A**, purple). Of all recorded PYR, 1,213 (22%) exhibited a consistent increase in firing rate during illumination (*p*<1.11×10^-16^, exact binomial test).

**Figure 5:**
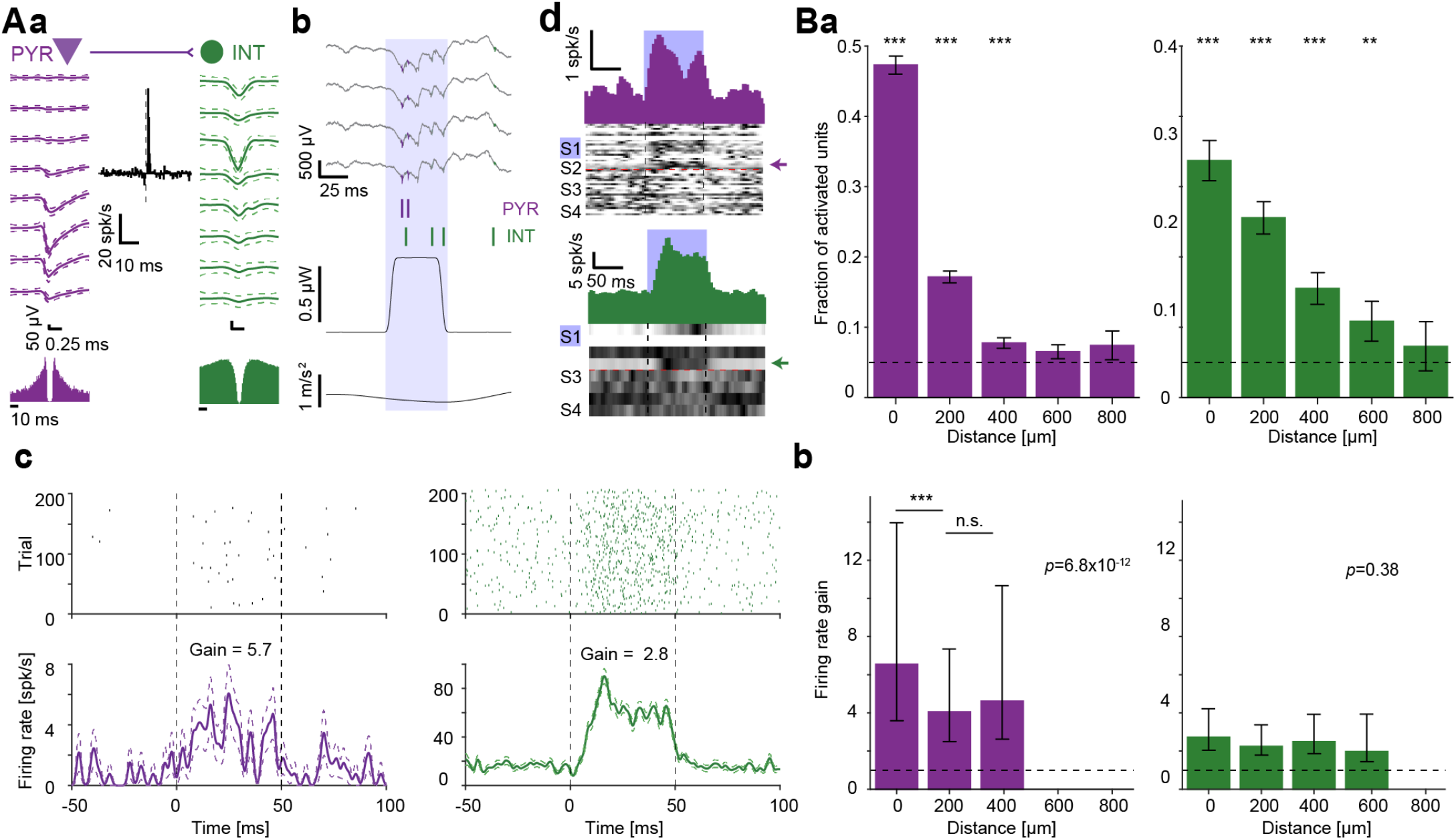
Transgenic hybrid mice enable optogenetic control of individual CA1 neurons. **(A)** Directly activated PYR and an indirectly activated INT recorded in hippocampal region CA1 of a HYB dual-transgenic (CaMKII::ChR2) mouse. (**a**) The units exhibit a cross-correlation histogram (*center*; no light condition) consistent with monosynaptic excitation (*p*<0.001, Poisson test). (**b**) Local field and spiking responses to a single 50 ms blue-light pulse (0.67 μW). The PYR and the INT both spike during the light. (**c**) Raster plots (*top*) and PSTHs (*bottom*) during blue light pulses for the PYR (*left*) and the INT (*right*). For visualization purposes, raster plots include 200 pulses. (**d**) *Top parts*: Mean PSTHs of 22 PYR (*purple*) and four INT (*green*) recorded simultaneously on the illuminated shank (S1). *Bottom parts*, *greyscale*: PSTHs of all 48 PYR and eight INT recorded simultaneously during the session. Each row shows the PSTH of one unit, scaled to the 0-1 (white-black) range for visualization purposes. Arrows indicate the example PYR and INT. **(B)** Activation probability of CA1 units depends on the distance from the illuminated shank. Dataset includes a total of 5,568 PYR and 1,066 INT. (**a**) Bars show the fraction of units with increased firing rate (*p*<0.05, Poisson test) during single-shank illumination. Error bars, SEM. ***: *p*<0.001, exact binomial test, comparing the fraction of light-activated units to chance level (horizontal dashed line). (**b**) Median firing rate gain of activated PYR and INT. Error bars, IQR. Gain is shown only for distances with an above-chance number of optically activated units. *P*-values indicated in text are for a one-way Kruskal-Wallis test. ***: *p*<0.001, Kruskal-Wallis test.

As previously reported in rat hippocampus (Stark et al., 2012), activation probability depended on the horizontal distance from the light source. Of the PYR recorded on the illuminated shank, 772/1,527 (47%) exhibited optical activation (*p*<1.11×10^-16^, binomial test), compared to 340/1,984 (17%) units recorded on the adjacent shank (horizontal spacing, 200 μm; *p*=2.5×10^-13^, binomial test; **Fig. 5Ba**, left). The fraction of optically-activated PYR was lower on shanks farther away (0 vs. 400 μm away: *p*<1.1×10^-16^, Bonferroni-corrected *G*-test). Moreover, the firing rate gain of the PYR that responded to focal illumination depended on the distance from the illuminated shank, being 6.5 [3.59 13.97] for same-shank PYR, compared to 4.1 [2.49 7.35] for PYR recorded on the adjacent shank (*p*=9.6×10^-10^, Kruskal-Wallis test; **Fig. 5Bb**, left). Therefore, CA1 PYR in dual-transgenic HYB exhibit robust distance-dependent responses to focal illumination.

Optogenetic stimulation led to the indirect activation of INT via monosynaptic inputs from excitatory neurons (**Fig. 5A**, green). A total of 241/1,066 (23%) of CA1 INT exhibited light-induced activation (*p*<1.11×10^-16^, binomial test; **Fig. 5Ba**, right). The fraction of light activated INT was 34% (92/273) on the illuminated shank and lower on shanks farther away (400 μm: 40/258, 16%, *p*=1.3×10^-4^, Bonferroni-corrected *G*-test). In contrast to PYR, the firing rate gain of light-activated INT did not differ consistently between shanks (*p*=0.38, Kruskal-Wallis test; **Fig. 5Bb**, right). Thus, consistent with previous observations in C57 mice (Stark et al., 2013), indirect INT activation in CA1 leads to more widespread activation than the local activation achieved in PYR.

To examine HYB optogenetic activation in a brain region with a distinct architecture, we performed optical stimulation in the neocortex of the same mice (n=21 sessions; **Table S5**). Similar to CA1 units, neocortical PYR and INT exhibited induced spiking (**Fig. S4A**), with activation probability that decreased with distance from the illuminated shank (**Fig. S4Ba**). A total of 274/409 (67%) neocortical PYR recorded on the illuminated shank exhibited optical activation, compared to 722/1,527 (47%) CA1 PYR (*p*=1.2×10^-4^, *G*-test). Furthermore, 331/591 (56%) neocortical PYR exhibited optical activation on the adjacent shank (200 μm away), compared to 340/1,984 (17%) CA1 PYR (*p*<3.5×10^-11^, *G*-test). Compared to CA1, focal illumination in the neocortex activated PYR located farther away from the illuminated shank. Light-induced firing rate gain did not differ between the illuminated shank (7.6 [4.54 16.03]) and the adjacent shank (7.6 [4.08 13.94]; *p*=0.79, Kruskal Wallis test; **Fig. S4Bb**, left). The difference between brain regions is consistent with different network topologies, since in contrast to the neocortex, CA1 PYR do not exhibit abundant recurrent excitation (Thomson and Radpour, 1991; Buhl et al., 2007). Neocortical units on distant shanks may be activated indirectly via excitatory synaptic connections from directly activated PYR closer to the illumination source.

An alternative approach for establishing optogenetic control in HYB is to cross FVB females with single-transgenic driver male mice and to inject the offspring with a viral vector that expresses a reporter gene. To demonstrate feasibility, we generated a PV-Cre x FVB mouse and injected the animal with a viral vector that allows Cre-dependent expression of ChR2. Focal illumination in the transgenic HYB induced direct PV activation and indirect PYR silencing (**Fig. S4C**). In summary, crossing transgenic C57 with FVB yields transgenic HYB offspring, suitable for optogenetic experiments.

## Discussion

Compared to the parental C57BL/6J and maternal FVB/NJ progenitor strains, the first-generation hybrid offspring (FVB/NJ x C57BL/6J)F1 exhibited reduced anxiety-like behavior, improved learning, and enhanced spatial long-term memory. Upon aging, motor endurance and balance performance of C57 and FVB mice deteriorated, whereas the sensorimotor abilities of hybrid mice were largely maintained. HYB displayed reduced anxiety-like behavior and improved spatial long-term memory upon aging. HYB were larger, ran faster, and performed more trials during electrophysiological experiments on the linear track. HYB and C57 exhibited similar CA1 place cell rate and phase coding. Finally, optogenetic manipulations were readily achieved in transgenic HYB.

Mice have emerged as a dominant model for biomedical research, mainly due to their small size, high reproductive rate, and genetic similarities to humans. The same properties facilitated the engineering of multiple transgenic lines. The maintenance of transgenic lines, particularly the generation of double- or multi-transgenic mice, is greatly facilitated by inbred mice (Silva et al., 1997). However, inbred mice are suboptimal for neurobehavioral studies due to idiosyncratic recessive traits. Furthermore, C57 exhibit age-dependent deterioration, limiting the duration of prolonged experiments. C57 mice are also smaller than other strains, which is a disadvantage for electrophysiological studies involving implanted headgear. Since headgear weight is limited by body size, the range of studied behaviors is limited by animal weight. A single well-defined outbreeding step produces larger offspring that exhibit enhanced behavior even during electrophysiology experiments, maintaining the ability of genetic targeting.

One of the challenges in mouse-based research is adjusting the strain to the specific study question. Strain selection can potentially affect research outcome, since strain characteristics may interact with genetic manipulations (Sittig et al., 2016). Furthermore, strain characteristics could interact with experimental manipulations. For example, mice that suffer from retinal degeneration cannot fully utilize visual information and exhibit impaired behavior in hippocampal-based navigation tasks that rely on visual cues (Nguyen et al., 2000). A second challenge is to ensure the stability of the tested phenotype under the null condition throughout the study period. In many cases, the effects of a given manipulation are studied over weeks to months (Prevot et al., 2019; Namdar et al., 2020; Rahn et al., 2021). Deterioration of control phenotypic behavior could confound potential manipulation-based inter-group differences. HYB exhibit particularly stable performance during aging, which is potentially useful for long-term studies.

To provide a quantitative description of the HYB, we supplemented the array of standard assays with gait assessment and home cage behavior. The gait of all strains was inconsistent with abnormalities seen in motor disorders (Preisig et al., 2016; Rahn et al., 2021), yet every strain exhibited distinct gait patterns that were largely preserved upon aging. In addition, we observed apparent inter-strain differences of daily conduct in the home cage, including distinct time division between sheltering, wheel running, and feeding. Together, the weight, daily conduct, and gait differences indicate that the spontaneous behavior of the (FVB/NJ x C57BL/6J)F1 hybrids is distinct from that of both progenitor strains.

There are some situations in which hybrid mice are not advantageous. First, compared to the C57 progenitors, HYB exhibited similar or improved performance in every assay used, with one exception – balance performance in the 3-month-old age group. HYB impaired balance may result from weight differences between the strains since HYB weigh more, and weight affects rotarod performance (Mao et al., 2015). Although HYB balance performance improved upon aging, hybrids may be unsuitable for some research topics. Second, some genes expressed by progenitors may be dominant; for instance, FVB contain a *Disc1* mutation which has been shown to cause cognitive impairment even when heterozygous (Koike et al., 2006; Ritchie and Clapcote, 2013). Nevertheless, we found that the potentially deleterious effects of dominant genes on HYB are smaller than the adverse effects of inbreeding. Third, because HYB mice are more robust, genetic knockout manipulations may be less likely to show a consistent phenotypic change (Sittig et al., 2016). Fourth, it is not straightforward to use hybrids if a homozygous allele is required or if knockout of both copies of a given gene is required (although see Silva et al., 1997).

For studies that do not involve genetic manipulations, HYB can be used as-is, providing improved performance while maintaining minimal inter-subject differences. Specifically, HYB can be implanted with high-density electrode arrays for neocortical and hippocampal recordings over multiple months. To combine transgenic control with enhanced behavior, we used two distinct strategies. One simple strategy involves using a homozygous parent of a C57-background, e.g., a driver line, yielding heterozygous offspring that present one transgenic allele. A second possibility is for one of the parents to be multi-transgenic, e.g., derived by crossing C57-based driver and reporter lines. Yet a third strategy is to use a C57-based driver line and backcross the reporter onto an FVB/NJ background. Due to the large body of transgenic mice developed on the C57 background, the number of possible combinations is very large, allowing to harness the behavioral potential of the hybrids for a wide range of studies.

We focused on first generation (F1) hybrids to minimize genetic differences between individuals. While further breeding is expected to increase inter-subject genetic variability, F2 hybrids will necessarily express recessive traits not expressed by the F1 generation. For instance, 25% of the F2 hybrids are expected to be blind due to a functional copy of the *Pde6b* gene (Pittler and Baehr, 1991) maintained in every F1 hybrid. Thus, unless careful backcrossing is performed, F2 hybrids should not be used. Second, we employed HYB which are offspring of FVB females and C57 males. Full characterization is required prior to the use of reciprocal hybrids, offspring of a C57 female and an FVB male. However, due to the reduced litter sizes of female C57 compared to female FVB (Taketo et al., 1991; Silver, 1995), the usage of the reciprocal hybrids is not recommended. Finally, since inter-strain differences are also sex-dependent (Brown and Wong, 2007), female HYB may exhibit distinct phenotypes.

## Conclusion

We showed that a single breeding step produces a hybrid mouse strain that exhibits improved behavioral performance while maintaining genetic control. The hybrid vigor and the enhanced behavioral capabilities may yield more trials in every experimental session, shorter learning periods, and higher accuracy in complex tasks. By increasing yield and accuracy, the usage of HYB may allow uncovering unexplored neuronal mechanisms. Enhanced behavior also has ethical advantages, since faster learning and higher yield per animal translate to fewer experimental animals or sessions required to answer a given question. The combination of genetic tools and the enhanced behavioral capabilities of the hybrid mice offers a unique opportunity for studying the neuronal basis of behavior.

## Acknowledgements

We thank Ronni Cohen for constructive comments. This work was supported by the United States-Israel Binational Science Foundation (BSF) 2015577; by the European Research Council 679253; by the Israel Science Foundation 638/16; by the Israel Science Foundation FIRST Program 1871/17; by the Rosetrees Trust A1576; and by the Canadian Institutes of Health Research (CIHR), the International Development Research Centre (IDRC), the Israel Science Foundation (ISF) and the Azrieli Foundation 2558/18.

## Author contributions

L.Sj. and E.S. conceived the project. L.B. and E.S. designed the experiments. H.E.S., A.L., O.A.V., T.K., L.Sp., S.So., R.G., and S.Si. bred and maintained animals. H.E.S. and E.S. implanted animals. H.E.S. and A.L. carried out electrophysiological experiments. H.E.S., L.B., A.L., and E.S. analyzed data. H.E.S., L.B., and E.S. wrote the manuscript, with contributions from all authors.

## Materials and Methods

### Experimental model and subject details

A total of 176 freely-moving adult mice were used in this study. 170 male mice were used for phenotyping, of which 61 were C57BL/6J (C57; JAX #000664, The Jackson Laboratory); 52 were FVB/NJ (FVB; JAX #001800); and 57 were hybrid (HYB; **Table S1**), offspring of an FVB female and a C57 male. Six transgenic mice were used for electrophysiological recordings (**Table S2**). One was single-transgenic and hybrid, generated by crossing an FVB female with a PV-Cre male (#008069). Two were dual-transgenic, generated by crossing CaMKII-Cre females (JAX #005359) with Ai32 males (#012569). One was dual-transgenic, generated by crossing an PV-Cre female with a Ai32 male. Two were dual-transgenic and hybrid, generated by crossing FVB females with second-generation (CaMKII-Cre x Ai32) males. All mice were bred in-house. After separation from the parents, animals were housed in groups of same-litter siblings. Animals were held on a reverse dark/light cycle (dark phase, from 8 AM until 8 PM). Data recorded from CA1 during linear track behavior were used in a previous report (Sloin et al., 2022). All animal handling procedures were in accordance with Directive 2010/63/EU of the European Parliament, complied with Israeli Animal Welfare Law (1994), and approved by the Tel Aviv University Institutional Animal Care and Use Committee (IACUC # 01-16-051, #01-19-017, and #01-21-051).

### Phenotyping study design

Mice of each strain were divided into two age groups, 3-month-old and 9-month-old (**Table S1**). Each age group was further divided into two subgroups: one subgroup was tested on test Battery A, and a second subgroup was used for test Battery B. Animals assigned to Battery A were subjected to assays in the following order: elevated plus maze (one day); open field (one day); rotarod (five days); treadmill (three days); optomotor drum (one day); and forced swim test (one day). Animals assigned to Battery B were tested on the catwalk (one day) and on the Morris water maze (five days). Mice were used in the phenotyping cages at least a week post-battery and were counterbalanced from Batteries A and B (9-month-old mice only). The division of assays into batteries was done to limit the number of assays that each mouse would perform. The order of assays in each battery was designed so that each battery began with voluntary, low stress eliciting assays. Between assays, mice were given a minimum of 24 hours without human interaction. All tests were initiated at beginning of the dark phase (8 AM) and were administered to a single mouse at a time. All equipment was thoroughly cleaned with Virusolve before and between trials. Except for the rotarod and treadmill tests, behavior during all tests was recorded using GigE cameras (ac1300-60gm mono, Basler; frame rate, 25 Hz). Commercial software (Ethovision 15XT; Noldus Information Technology, Wageningen, The Netherlands; Noldus et al., 2001) was used to analyze all video files.

#### Elevated plus maze

The elevated plus maze is used to evaluate anxiety-like behavior based on the natural uneasiness of rodents towards open, elevated fields (Rodgers and Dalvi, 1997; Crawley, 2000; Komada et al., 2008). The apparatus consists of a four-armed platform resembling a shape, positioned 40 cm above the floor. Two arms are confined by walls (“closed”; L×W×H: 35×5×15 cm), whereas two other arms are not walled (“open”; 35×5 cm). Similar arms face one another. In the test, the mouse was initially placed at the center of the maze facing one of the closed arms and then allowed to move freely for 7 min. The time spent exploring the open arms (“open”) and the time spent exploring the closed arms (“closed”) were used to derive an index (open-closed)/(open+closed) that served as a contraindication to anxiety-like behavior (“contra-anxiety index”).

#### Forced swim test

The forced swim test is used to evaluate depressive-like behavior based on induced “behavioral despair” (Porsolt et al., 1977; Castagné et al., 2010; Can et al., 2012). The apparatus consists of a clear plexiglass cylinder (24 cm in height, 19 cm diameter) filled with 16 cm of water at 22 °C. In the test, the mouse was placed in the cylinder for 7 min and then moved to a heated cage until the fur dried completely. Typically, the mouse gradually stopped swimming before being removed. Freeze (immobility) duration, defined as the time the mouse remained floating motionless in the water, was measured during the last 5 min as a manifestation of behavioral despair.

#### Gait analysis

The gait analysis apparatus (CatWalk XT, Noldus Information Technology, Wageningen, The Netherlands) enables the assessment of voluntary gait and locomotion in mice (Moller et al., 2000; Crowley et al., 2018). The apparatus consists of a hardware system with a glass walkway (L×W: 130×20 cm), a GigE video camera, and a software package for the quantitative assessment of animal footprints. The walkway is illuminated from above by red light and from the side by green light. The green light is internally reflected within the glass, except at touched points. The walkway is connected to a dark goal box and enclosed by an adjustable tunnel at one end. In the test, mice were placed at the walkway entrance and allowed to run freely. A successful run was defined when the animal traversed the track without pausing. Every animal was tested until three successful runs were completed. For every parameter, the average of the three runs was used for analysis.

#### Morris water maze

The Morris water maze (MWM) is used to evaluate learning and spatial long-term memory based on the natural tendency of rodents to attempt to escape a body of water (Morris, 1981; Morris, 1984; Vorhees and Williams, 2006; Bromley-Brits et al., 2011). The apparatus is a circular pool (150 cm diameter, 10 cm depth) filled with water and maintained at a temperature of 23 °C. A transparent platform (5×5 cm) was fixed 1 cm below the water surface in a constant location. The pool was situated in a room containing distal visual cues, and proximal cues were placed on the inner walls of the pool. The mouse was introduced to the pool at different starting points and allowed 60 s to find the platform and an additional 15 s to stay on the platform. If the mouse failed to find the platform within 60 s, the animal was guided to the platform and allowed to stay there for 15 s. The animal swam three times every day during four consecutive “learning days”. During the learning days, the time before finding the platform (“latency to platform”) and the total distance traveled were measured. Changes between days were used to evaluate learning performance.

On the fifth day of the assay (“probe day”), the platform was removed, and the mouse was allowed to swim for 60 s. The time spent in every quadrant was recorded. The fraction of time spent in the “target” quadrant (i.e., close to the location of the missing platform), the number of visits to the target quadrant, and the latency to platform were used to evaluate spatial long-term memory performance.

#### Open field

The open field test provides a way to systematically assess motivational behavior by quantifying exploration of a novel environment and general locomotor activity (Christmas and Maxwell, 1970; Prut and Belzung, 2003). In the test, the mouse was initially placed in one of the corners of an open-top plexiglass box (L×W: 50×50 cm, raised 40 cm above the floor), and behavior was recorded for 15 min. Activity, defined as the fraction of pixels that changed in the entire arena between consecutive frames (frame rate, 25 Hz), was measured throughout the run duration to evaluate exploratory behavior.

#### Optomotor visual test

The optomotor drum is based on an apparatus developed for immobile mice (Michener et al., 1979) and allows assessing visual behavior in freely-moving mice (Abdeljalil et al., 2005). The apparatus consists of an elevated stationary platform (20 cm) surrounded by a drum (internal diameter, 39 cm) with vertically oriented black and white stripes on the inside, each spanning 10°. In the test, the mouse was initially habituated to the platform for 2 min, with the drum stationary. The drum then rotated at 2 rpm counterclockwise for 2 min, stopped for 30 s, and rotated clockwise for 2 min. The number of head turns (15° movements at the drum speed) and the cumulative duration of head turns were measured.

#### Phenotyping cages

The PhenoTyper (model 3000, Noldus Information Technology, Wageningen, The Netherlands) is an instrumented home cage in which rodent behavior is automatically monitored through a video- and event-based system (De Visser et al., 2006; Maroteaux et al., 2012; Grieco et al., 2021). The cage (L×W×H: 30×30×35 cm) is made of transparent plexiglass walls with an opaque plexiglass floor. The cage is equipped with a watering station, a feeding station, a running wheel, and a shelter in one corner (H×D: 10×9 cm, transparent material). The feeding and watering stations are equipped with beam-breaking devices, allowing automatic recording of feeding behavior and water intake (number of feeds and licks). The lid of the cage lid is equipped with an infrared-sensitive video camera (768 x 576 pixels) and several infrared LEDs, allowing continuous recording of animal position. Bedding covered the floor, and food and water were provided ad libitum. The mouse was introduced to the cage at the beginning of the dark phase (8 AM). Testing lasted three days, during which no human interference took place.

#### Rotarod test

Mice were subjected to a five-lane accelerating rotarod (Ugo Basile, Gemonio, Italy) to evaluate motor learning and balance (Jones and Roberts, 1968; Crawley, 2000; Crawley, 2003; Bailey et al., 2009; Shiotsuki et al., 2010). The apparatus consists of a 3.2 cm (diameter) horizontal rod elevated 10 cm from the ground. The paradigm was comprised of five consecutive days. On every day, mice were subjected to five trials, of which the duration of the three longest trials were averaged. A trial began with the rod rotating at 4 rpm and gradually accelerating for 5 minutes, up to a maximum of 50 rpm. The duration until the animal fell from the rod (“latency to fall”) was measured.

#### Treadmill

The treadmill apparatus (Panlab, Harvard Apparatus, Barcelona, Spain) is used to assess maximal endurance in mice (Marques-Aleixo et al., 2015). The apparatus consists of a five-lane motorized treadmill with an electric shock zone at one end of the track. Each lane is 38×7×7 cm (L×W×H). A current shock (0.2 mA) was used to encourage running; cumulative shock duration of 2 s defined the maximal ability of mice to run. The protocol included two training days and one testing day. On the first training day, treadmill speed was fixed at 5 cm/s. The second training day consisted of two parts: (i) walking on the treadmill for 300 s at a constant speed (5 cm/s); (ii) subjecting the mice to 330 s of locomotion at a variable speed (increasing from 5 to 21 cm/s by 1 cm/s every 20 s). On the third (testing) day, speed was initially set to 5 cm/s and was gradually increased by 1 cm/s every 20 s. The trial ended when the cumulative shock duration was 2 s. Trial duration and distance to failure were measured.

### Indices for behavioral assays

An “aging index” was defined for every parameter as the difference between the median value of that parameter for 9-month-old mice and the median for 3-month-old mice of the same strain, divided by the sum. We used a bootstrap procedure to estimate index dispersion and determine statistical significance. The null hypothesis of the index being equal to zero is equivalent to no consistent changes between the two age groups. In the procedure, we resampled the data (e.g., 12 values from 3-month-old mice and 15 values from 9-month-old mice) with replacement many (10,000) times. For each resampling iteration, we computed the aging index. The reported index is the mean over all iterations, and the SEM is the SD over all iterations. For a one-sided alternative hypothesis (increase/decrease), *p*-values are the fraction of indices below/above zero.

A “C57 index” was defined in an equivalent manner, where the index was defined as the difference between the median value of the parameter of interest for the tested strain (FVB or HYB) and the median value of the same parameter for C57 mice of the same age group, divided by the sum. Mean, SEM, and *p*-values were estimated using bootstrapping as for the aging indices.

When parameters are strictly positive, the difference divided by the sum, (a-b)/(a+b), provides a valid estimate. However, some parameters of interest can take negative values: for instance, the “contra-anxiety index” ranges from −1 to 1. Then, an increase from one negative value to another will yield a negative “aging index” (or “C57 index”). Thus, when the parameter of interest was itself an index, we replaced the ratio (a-b)/(a+b) with the a/b ratio.

### Probes and surgery

Every animal used in electrophysiological experiments (**Table S3**) was implanted with a multi-shank silicon probe attached to a movable microdrive, equipped with optical fibers following previously described procedures (Stark et al., 2012; Noked et al., 2021). The probes used were Stark64 (Diagnostic Biochips), Buzaski32 (NeuroNexus), and Dual-sided64 (Diagnostic Biochips). The Stark64 probe consists of six shanks, spaced horizontally 200 μm apart, with each shank consisting of 10-11 recording sites spaced vertically 15 μm apart. The Buzaski32 probe consists of four shanks, spaced horizontally 200 μm apart, with each shank consisting of eight recording sites spaced vertically 20 μm apart. The Dual-sided64 probe consists of two dual-sided shanks, spaced horizontally 250 μm apart, with each shank consisting of 16 channels on each side (front and back), spaced vertically 20 μm apart.

Prior to probe implantation, the single-transgenic hybrid was injected with a DIO-hChR2 viral vector (rAAV5/EF1a-DIO-hChR2(H134R)-eYFP; 3.2×10^12^ IU/mL; University of North Carolina viral core facility; courtesy of K. Deisseroth). The solution was injected stereotactically (Kopf) into the neocortex and hippocampus at 8 different depths (AP −1.6, ML 1.1, DV 0.4 to 1.8 at 0.2 mm increments; 25 nl/site; Nanoject III, Drummond).

Probes were implanted in the parietal neocortex above the right hippocampus (AP/ML, −1.6/1.1 mm; 45° angle to the midline) under isoflurane (1%) anesthesia. Following recovery from anesthesia, linear-track animals were placed on a water-restriction schedule that guaranteed at least 40 ml/kg of water (corresponding to 1 ml per 25 g mouse) on every recording day. Recordings were carried out five days a week, and animals received free water on the sixth day. After every one to five recording sessions, the probe was translated vertically downwards by up to 70 μm.

### Linear track sessions

For the linear track analyses, neuronal activity was recorded in 4.5 [1.9 10.1] hour sessions (median [IQR]). At the beginning of every session, neural activity was recorded while the animal was in the home cage. The animal was then placed on a 150 cm linear track that extended between two 10 x 10 cm square platforms. Each platform included a water delivery port. Mice were under water restriction and were trained to repeatedly traverse the track for a water reward of 3-10 μL. Over all sessions, mice ran 167 [132 200] one-direction trials over about one hour (**Table S4**). Trials with a mean running speed below 10 cm/s were excluded from analyses. Animals were equipped with a 3-axis accelerometer (ADXL-335, Analog Devices) for monitoring head movements. Head position and orientation were tracked in real-time using two headmounted LEDs, a machine vision camera (ace 1300-1200uc, Basler), and a dedicated system (“Spotter”, Gaspar et al., 2019).

### Spike detection and sorting

Neural activity was filtered, amplified, multiplexed, and digitized on the headstage (0.1–7,500 Hz, x192; 16 bits, 20 kHz; RHD2132 or RHD2164, Intan Technologies) and then recorded by an RHD2000 evaluation board (Intan Technologies). Offline, spikes were automatically detected and sorted into single units using KlustaKwik3 (Kadir et al., 2014; Rossant et al., 2016) for shanks with up to 11 sites/shank or KiloSort2 (Pachitariu et al., 2016) for 16 channel shanks. Automatic spike sorting was followed by manual adjustment of the clusters. Only well-isolated units were used for further analyses (amplitude >40 μV; L-ratio <0.05 (Schmitzer-Torbert et al., 2005); ISI index <0.2 (Fee et al., 1996)). Units were classified into putative PYR or PV-like INT using a Gaussian mixture model (Stark et al., 2013).

### Place field and phase precession analysis

Based on the linear track data, spatial information, place fields, and phase precession were determined for every PYR (Sloin et al., 2022). Briefly, place fields were defined as regions spanning 15-100 cm in which the firing rate increased compared to the on-track spontaneous firing rate (p<0.05, Bonferroni-corrected Poisson test). Theta phase precession was quantified for each place field using a circular-linear analysis (Schmidt et al., 2009; Kempter et al., 2012). The circular-linear model yielded the precession slope, *a*, and the resultant length of the residuals, *R*, indicating model fit; statistical significance was determined by a permutation test (Sloin et al., 2022). Precession effect size was quantified as the ratio between the fit of spikes to the circular-linear model, *R*, divided by the median of 300 model fits to randomly permuted phase/position pairs.

### Optogenetic stimulation

To determine the effect of optogenetic stimulation on spiking, 50 ms blue-light pulses were administered. In every session, illumination was carried out for every shank separately. In CA1 (**Fig. 4**), pulses were given in a total of 131 shanks during 55 sessions, using light power of 2.43 [0.96 3.98] μW. In a given stimulation experiment, pulses were applied 137 [51 209] times. In the neocortex (**Fig. S4**), pulses were given in a total of 63 shanks during 21 sessions, using light power of 11.07 [5.25 21.71] μW. In a given stimulation experiment, pulses were applied 150 [75 300] times. Light-induced firing rate gain was defined as the mean firing rate during illumination, divided by the mean firing rate during baseline (in the lack of illumination on any shank). Units were determined as light-activated if the Poisson probability of seeing the observed number of spikes (or more) during illumination was *p*<0.05, based on the baseline firing.

### Statistical analyses

In all statistical tests used in this study, a significance threshold of α=0.05 was used. All descriptive statistics (n, median, IQR, mean, SEM) can be found in the results, the figure legends, and the tables. Differences between medians of two groups were tested with Mann-Whitney’s *U*-test. Differences between medians of three groups or more were tested with Kruskal-Wallis nonparametric analysis of variance and corrected for multiple comparisons using Tukey’s procedure. Differences between medians measured along two dimensions (e.g., strain and day) were tested with a two-way Kruskal-Wallis analysis of variance. Wilcoxon’s signed-rank test was employed to determine whether a group median is distinct from zero. To estimate whether a given fraction was smaller or larger than expected by chance, an exact binomial test was used. Differences between the proportions of observations of two categorical variables were tested with a likelihood ratio (*G*-) test of independence. Bonferroni’s correction was employed in cases of *G*-test multiple comparisons.

## Supplementary material for

**Figure S1:**
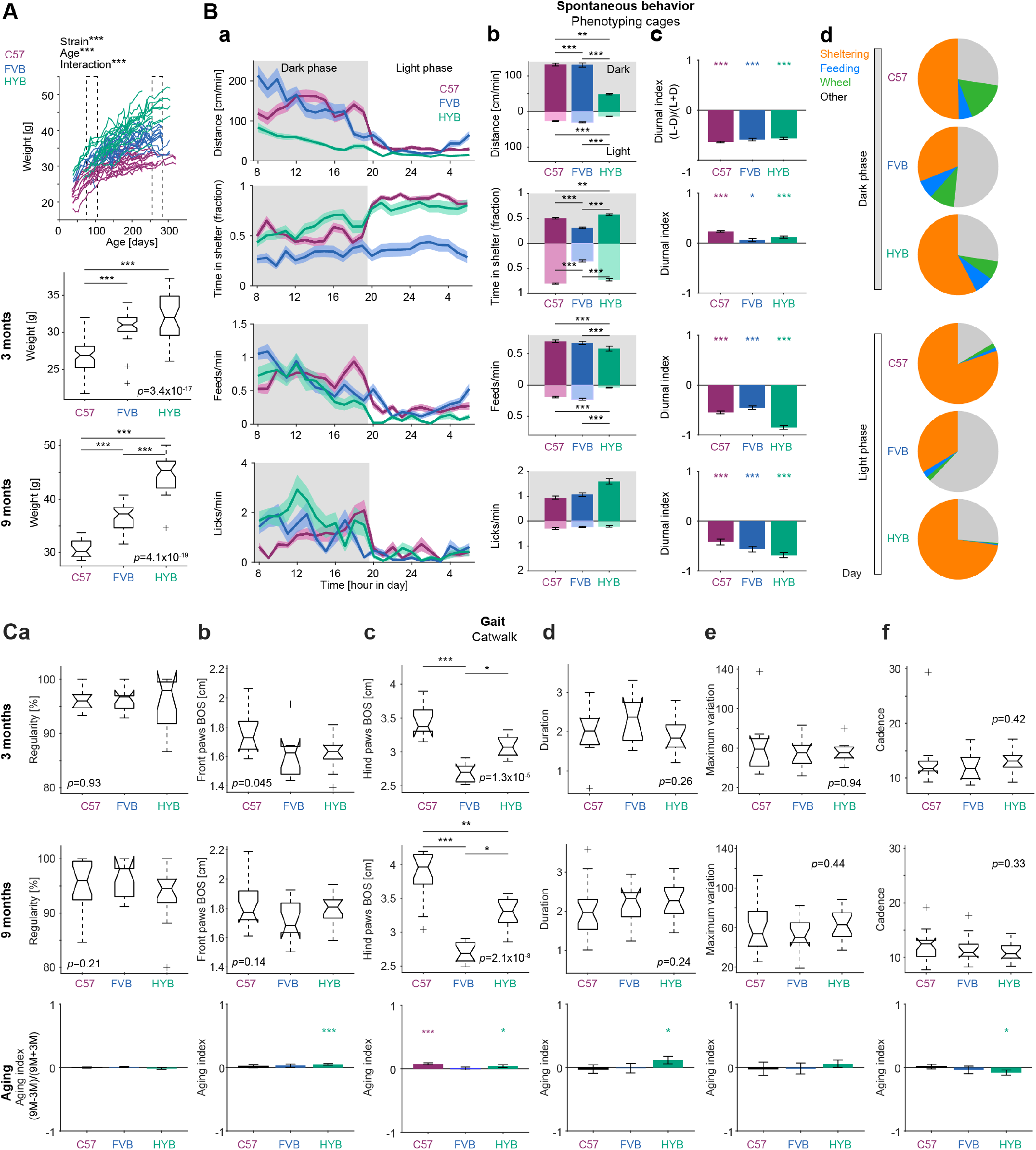
Hybrids are larger and exhibit distinct patterns of daily conduct in the home cage. **(A)** HYB weigh more than both progenitors. *Top*: Mice of the three strains were weighed periodically over 6 months, from 3-month-old to 9-month-old (n=20 C57, 17 FVB, and 15 HYB). Each line connects measurements of a single animal. ***: *p*<0.001, strain by day two-way Kruskal-Wallis test. Dashed lines, age ranges used for statistical analyses (*center* and *bottom* panels). *Center and bottom*: Every box plot shows median and inter-quartile range (IQR), whiskers extend for 1.5 times the IQR in every direction, and a plus indicates an outlier. For every mouse, weights were averaged over measurements made in the age of 75-105 days (“3 months”; *center*) and 255-285 days (“9-months”; *bottom*). *P*-values indicated by text are for one-way Kruskal-Wallis test. ***: *p*<0.001, Kruskal-Wallis test, corrected for multiple comparisons. **(B)** Hybrids are least active and spend more time in the shelter during the dark phase. 9-month-old mice (n=16 C57, 15 FVB, and 11 HYB) were placed in individual phenotyping cages and the distance covered, time in the shelter, feeding, and licking were sampled every hour over three days. (**a**) *Top*: During both dark (0800 until 2000) and light phases, HYB cover less distance. Bands show SEM. *Second row*: During the dark phase, HYB spend more time in the shelter than C57 or FVB. *Third row*: During both dark and light phases, HYB feed less than either C57 or FVB. *Bottom*: The number of licks is not consistently different between the strains. (**b**) Mean metric (distance covered, sheltering, feeding/licking rate) during the dark (*top half*) and light (*bottom*) phases. Error bars, SEM; **/***: *p*<0.01/*p*<0.001, Kruskal-Wallis test, corrected for multiple comparisons. (**c**) Diurnal indices, defined as the difference between the metric (e.g., distance covered) during the light phase (L) and the dark phase (D), divided by the sum. */***: *p*<0.05/*p*<0.001, bootstrap test. Mice of all strains exhibit diurnal changes in all metrics tested. (**d**) Pie charts show the fraction of time spent by mice of every strain carrying three specific behaviors: sheltering (being in the shelter), feeding (breaking the photobeam of the feeding station), and wheeling (being on the wheel). “Other” corresponds to home cage roaming, including licking. The time mice dedicated to each behavior differs between strains (*p*<0.001, strain, behavior, and interaction effects, two-way Kruskal-Wallis test). Compared to the dark phase, mice of all strains spend less time on the wheel and feeding during the light phase (*p*<0.001, Kruskal-Wallis test). **(C)** Gait patterns of the three strains. (**a**) Gait regularity does not differ consistently between strains or between age groups. Here and in all other panels, *p*-values indicated in text in the *top* and *center* are for Kruskal-Wallis test (e.g., 0.93). *Bottom*: Aging indices. Bars, mean aging index; error bars, SEM; */***: *p*<0.05/*p*<0.001 bootstrap test, compared to a no-change null. (**b**) The front paws base of support (BOS) of HYB increases with aging. (**c**) The hind paws BOS is distinct for the three strains. */**/***: *p*<0.05/*p*<0.01/*p*<0.001, Kruskal-Wallis test, corrected for multiple comparisons. (**d, e, f**) Track crossing duration, maximum variation, and cadence do not differ consistently between strains in either age group.

**Figure S2:**
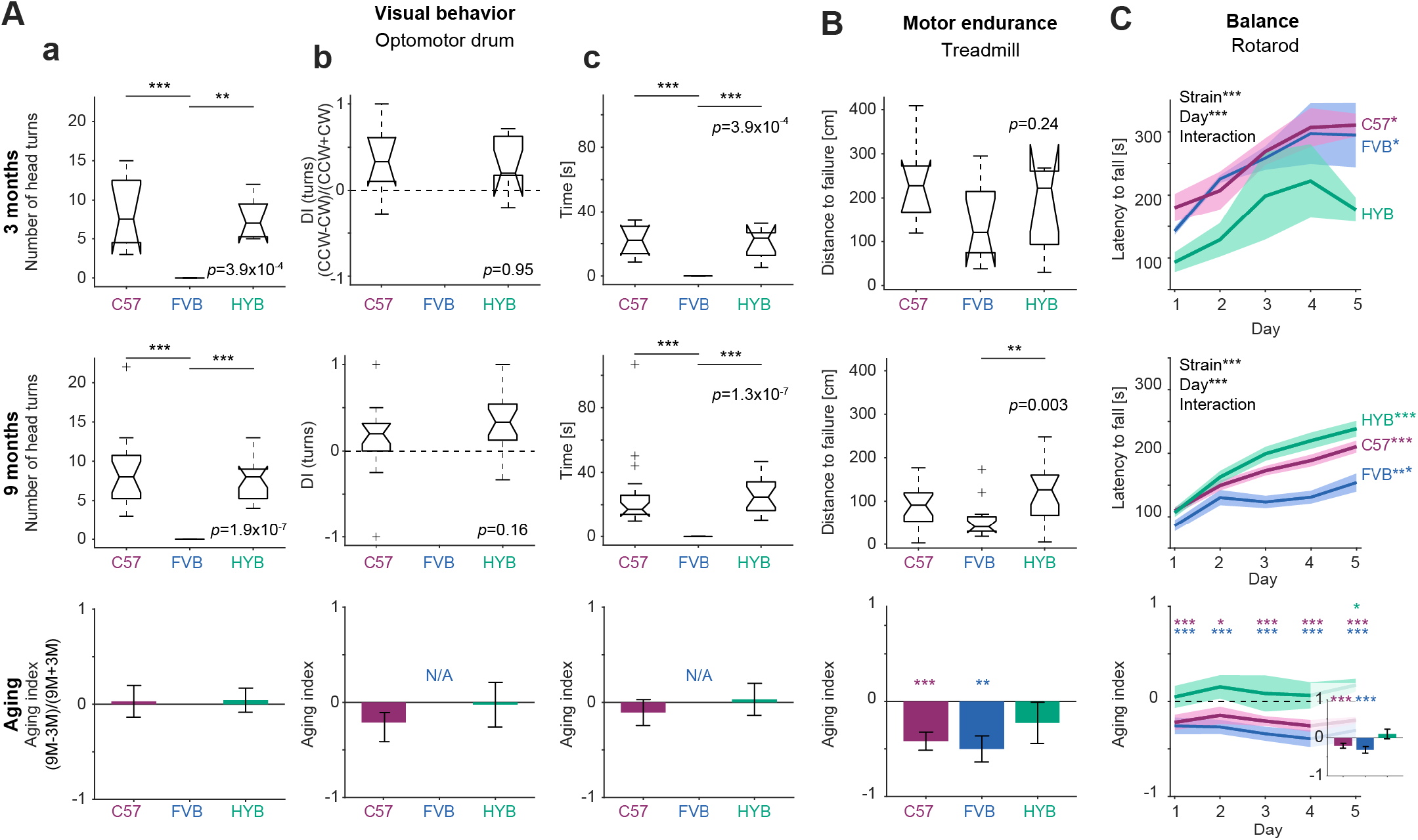
Hybrid sensorimotor performance is maintained across age groups. **(A)** Reaction to visual stimuli on the optomotor drum of HYB and C57 is similar and remains intact at an older age. In contrast, FVB do not react to rotating visual stimuli at either age group. *Top row*: 3-month-old mice; *center*: 9-month-old mice; *bottom*: aging indices (**a**) FVB mice do not make any head turns, indicating poor visual abilities. Here and in other panels, *P*-values indicated in text are for a Kruskal-Wallis test. **/***: *p*<0.01/*p*<0.001, Kruskal-Wallis test, corrected for multiple comparisons. (**b**) Directional bias in the optomotor drum. Every trial included 2 min of 2 rpm counterclockwise (CCW) and 2 min of 2 rpm clockwise (CW) drum rotations. Directionality indices (DI) were defined as the number of head turns during CCW vs. CW drum rotations. (**c**) Time spent making head movements. FVB mice do not move their heads as the drum turns, whereas the time spent moving the heads does not differ consistently between C57 and HYB in either age group. **(B)** Motor endurance deteriorates for older C57 and FVB mice. *Center*: At the age of 9 months, HYB cover longer distances than FVB. **: *p*<0.01, Kruskal-Wallis test, corrected for multiple comparisons. *Bottom*: **/***: *p*<0.01/*p*<0.001, bootstrap test. In contrast to C57 and FVB, HYB performance is maintained between age groups. **(C)** Balance performance deteriorates upon aging for C57 and FVB. In contrast, HYB balance performance is maintained between age groups. *Top* and *center*: Mean (SEM) lantecy to fall on five consecutive training days; *** next to the text indicate *p*<0.001 for a two-way, strain by day, Kruskal-Wallis test. */*** at right: *p*<0.05/p<0.001, one-way Kruskal-Wallis test, measuring within-strain, across-day effect. In the older age group, HYB remained on the apparatus for the longest duration before falling. *Bottom*: Aging indices were computed for each strain every day. */***: *p*<0.05/p<0.001, bootstrap test. *Inset*: Aging indices, averaged over days.

**Figure S3:**
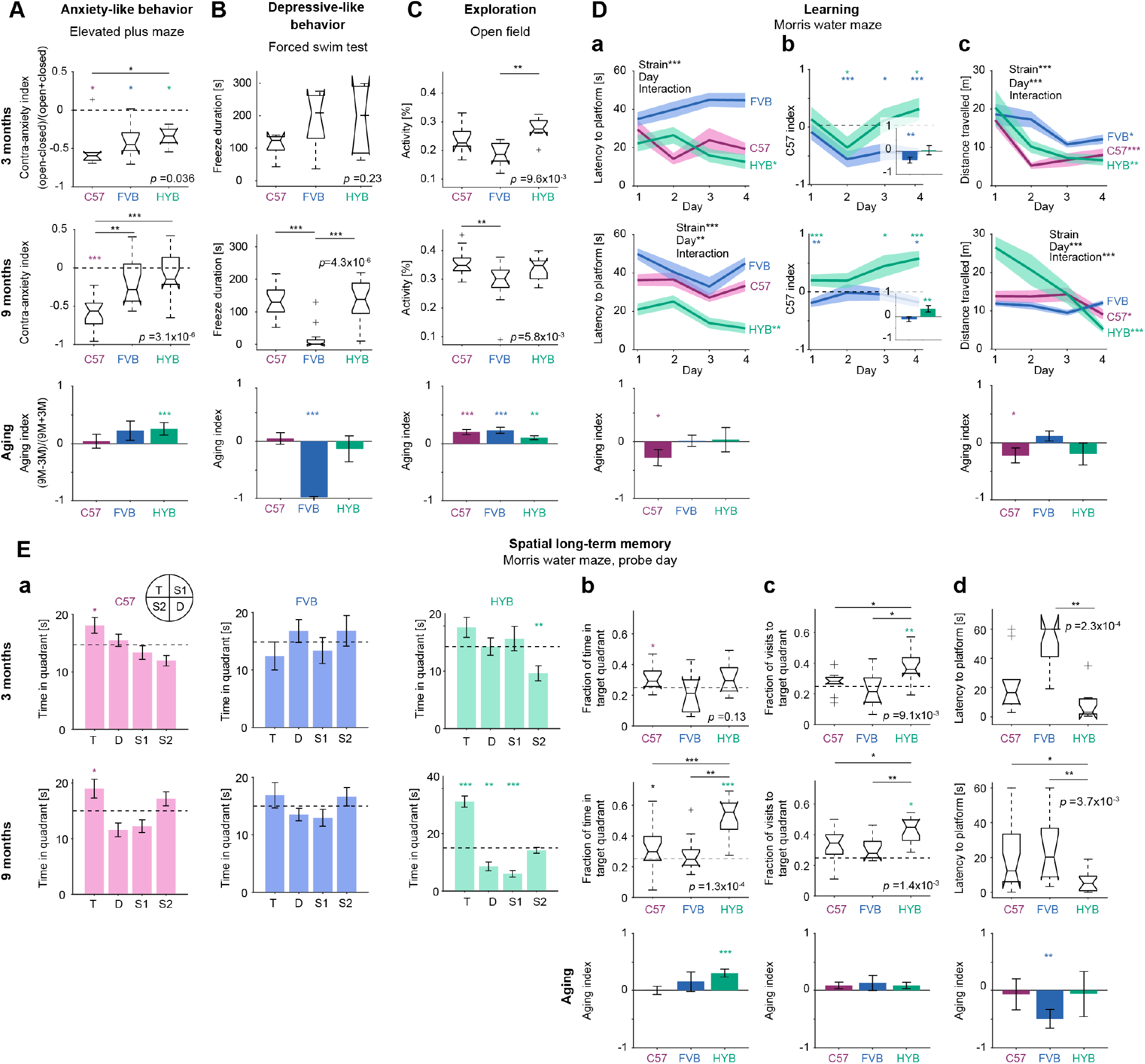
Hybrids exhibit reduced anxiety-like behavior, improved learning, and improved memory. **(A)** HYB explore the open arms of an elevated plus maze more than C57 mice, particularly at an older age, indicating reduced anxiety-like behavior. Here and in subsequent panels: *Top* and *center*: *P*-values indicated by text are for a one-way Kruskal-Wallis test; lined */**/***: *p*<0.05/*p*<0.01/*p*<0.001, Kruskal-Wallis test, corrected for multiple comparisons. */**/***: *p*<0.05/*p*<0.01/*p*<0.001, Wilcoxon’s signed-rank test, comparing indices to chance level. *Bottom*: Aging index. Bars and errors show mean and SEM, respectively; */**/***: *p*<0.05/*p*<0.01/*p*<0.001, bootstrap test, compared to a no-change null (index of zero). **(B)** Depressive-like behavior is not consistently different between HYB and C57 mice, and is reduced for older FVB mice. **(C)** Open field exploration increases at an older age for all strains. *Top*: In the 3-month-old age group, HYB are more active in the open field than FVB. *Center*: In the 9-month-old group, C57 were more active than FVB. *Bottom*: Mice of all three strains exhibit increased activity upon aging, evident as above-zero aging indices. **(D)** Learning on the Morris water maze (MWM) task is improved for HYB compared to C57, and is maintained at older age. (**a**) Mean (and SEM) latency to finding the platform (limited to 60 s). Here and in **c**: **/*** next to the text indicate *p*<0.01/*p*<0.001 in a two-way, strain by day, Kruskal-Wallis test; */**/*** at right indicate *p*<0.05/*p*<0.01/*p*<0.001, one-way Kruskal-Wallis test, measuring the within-strain, across-day effect. In both age groups, only HYB show consistent across-day learning. *Bottom*: Aging indices, averaged over all learning days. Learning performance of C57 deteriorates at an older age. (**b**) Day-by-day “C57 indices” (**Fig. 2D**), measuring the change in the latency to platform between FVB (blue) or HYB (green) and C57 mice. Bands show mean and SEM. */**/***: *p*<0.05/*p*<0.01/*p*<0.001, bootstrap test. *Insets*: mean (and SEM) C57 indices, averaged over all learning days; *p*-values, geometric mean over all learning days. (**c**) C57 and HYB travel progressively shorter distances over training days, indicating learning. All conventions are the same as in **a**. **(E)** On the probe day of the MWM task, HYB spend the longest time in and the shortest latency to the target quadrant. (**a**) Time spent in every quadrant. */**/***: *p*<0.05/*p*<0.01/*p*<0.001, Wilcoxon’s signed-rank test, comparing time in each quadrant to chance level (0.25 of the total time). (**b**) Time spent in the target quadrant, for 3-month-old (*top*) and 9-month-old (*center*) mice. Here and in **c**: Horizontal dashed lines indicate chance level of 0.25, expected for random exploration; other conventions are the same as in **A**. Compared to C57 or FVB, 9-month-old HYB spend a larger fraction of time in the target quadrant. *Bottom*: Aging indices. Compared to younger HYB, older HYB spend more time in the target quadrant. (**c**) In both age groups, the fraction of visits to the target quadrant is highest for HYB. (**d**) Latency to the platform location, measured during the probe day, is the shortest for HYB.

**Figure S4:**
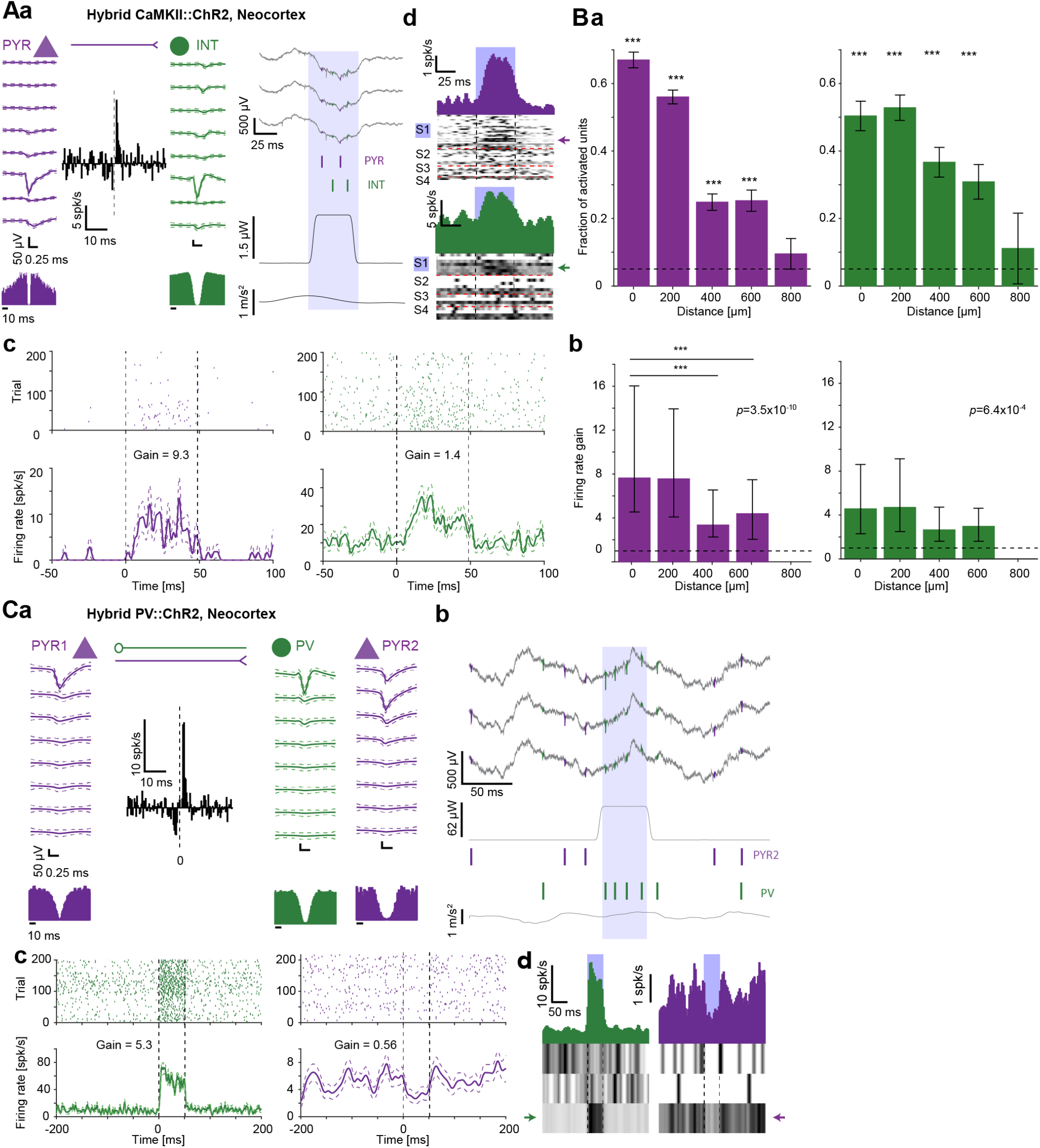
Transgenic hybrids support optogenetic control of individual neocortical neurons. **(A-B)** Optogenetic activation in the neocortex of hybrid freely-moving dual-transgenic CaMKII::ChR2 mice. **(A)** Example directly activated PYR and indirectly activated INT recorded in HYB neocortex. (**a**) The units exhibit a cross-correlation histogram (*center*; no light condition) consistent with monosynaptic excitation (*p*<0.001, Poisson test). *Bottom*: Auto-correlation histograms (ACHs). Wide-band (0.1-7,500 Hz) spike waveforms (mean±SD) recorded on the same shank; inter-site vertical spacing is 20 μm. (**b**) Local field and spiking responses of the units to a single 50 ms blue-light pulse (1.71 μW). Both units spike during the light. (**c**) Raster plots (*top*) and PSTHs (*bottom*) during 230 blue light pulses for the PYR (*left*) and indirectly activated INT (*right*). The raster plot shows 200 representative repetitions. (**d**) *Top*: Mean PSTHs of 26 PYR (*purple*) and seven INT (*green*) recorded simultaneously on the illuminated shank during light pulses. *Bottom*, *greyscale*: PSTHs of all units recorded during the session (49 PYR and 21 INT). S1, S2, S3, and S4 denote different shanks. Each row corresponds to the PSTH of one unit, scaled to the 0-1 (white-black) range for visualization purposes. Arrows indicate the example units. **(B)** Activation probability and firing rate gain of neocortical units depend on the distance from the illuminated shank. Dataset includes a total of 1,546 PYR and 517 INT. (**a**) Bars show the fraction of units with increased firing rate (*p*<0.05, Poisson test) during single-shank illumination as a function of the horizontal distance from the illuminated shank. Error bars, SEM. ***: *p*<0.001, exact binomial test, compared to chance level (0.05, horizontal dashed line). Neocortical PYR and INT recorded up to 600 *μ*m away from the illuminated shank are activated. (**b**) Median firing rate gain of activated PYR and INT as a function of distance from the illuminated shank. Error bars, IQR. Gain is considered only for distances with an above-chance number of activated units. *P*-values indicated in text are for a one-way Kruskal-Wallis test. ***: *p*<0.001, Kruskal-Wallis test, corrected for multiple comparisons. Gain differs between activated PYR recorded on the illuminated shank and activated PYR recorded 400 *μ*m away. **(C)** Parvalbumin-immunoreactive (PV) cell activation in a hybrid freely-moving single-transgenic PV-Cre mouse, injected with a Cre-dependent viral vector for ChR2. (**a**) Subnetwork of a PYR (PYR1; *purple*) and a PV cell (*green*), recorded simultaneously using a high-density optoelectronic probe. ACHs and cross-correlation histogram (no light condition), consistent with reciprocal monosynaptic excitation and inhibition (*p*<0.001, Poisson test). (**b**) Wide-band (0.1-7,500 Hz) traces and spikes of the PV cell and another unit, PYR2. The top three traces are neuronal recordings; the fourth trace indicates instantaneous light power emitted by the fiber attached to the recording shank; and the bottom trace indicates head acceleration. During blue light pulses (62 *μ*W), the PV cell spikes, whereas PYR2 spiking is suppressed. (**c**) Raster plot (*top*) and PSTH (*bottom*) during blue light pulses for the light-activated PV cell (*left*) and the suppressed PYR (*right*). Arrows indicate the example units. (**d**) PSTHs of six units recorded by the illuminated shank during blue light pulses. *Bottom*: Each row corresponds to the PSTH of one unit, scaled to the 0-1 (white-black) range for visualization purposes. *Top*: Mean PSTHs.

**Table S1:** Mice used for phenotyping assays.

|  | C57, 3M | FVB, 3M | HYB, 3M | C57, 9M | FVB, 9M | HYB, 9M | Total |
| --- | --- | --- | --- | --- | --- | --- | --- |
| **Battery A** | 12 | 12 | 11 | 19 | 14 | 19 | 87 |
| Elevated plus maze | 8 | 8 | 7 | 19 | 14 | 19 |  |
| Open field | 7 | 8 | 7 | 19 | 14 | 15 |  |
| Rotarod<sup>a</sup> | 4 | 4 | 4 | 19 | 14 | 19 |  |
| Treadmill | 8 | 8 | 7 | 19 | 14 | 19 |  |
| Optomotor drum | 8 | 8 | 7 | 19 | 14 | 19 |  |
| Forced swim test | 6 | 8 | 7 | 19 | 14 | 19 |  |
| Phenotyping cages<sup>b</sup> | - | - | - | 8 | 8 | 6 |  |
| **Battery B** | 10 | 9 | 12 | 20 | 17 | 15 | 83 |
| Weight<sup>c</sup> | - | - | - | 20 | 17 | 15 |  |
| Gait analysis | 10 | 9 | 11 | 19 | 12 | 15 |  |
| Morris water maze | 10 | 9 | 11 | 18 | 11 | 15 |  |
| Phenotyping cages<sup>b</sup> | - | - | - | 8 | 7 | 5 |  |
| **Total** | 22 | 21 | 23 | 39 | 31 | 34 | 170 |

<sup>a</sup>Due to errors in the initial testing of 3-month-old mice on the rotarod, an additional replacement cohort of 12 animals was used.
<sup>b</sup>Mice tested in the phenotyping cages were a subset of the animals used in Batteries A and B.
<sup>c</sup>Mice were weighed at least once every week from the age of 3 months until subjected to test Battery B at 9 months of age.

**Table S2:** Results of phenotyping assays for every age group and strain.

| Phenotype | Test | Metric | C57, 3M<sup>a</sup> | FVB, 3M<sup>a</sup> | HYB, 3M<sup>a</sup> | C57, 9M<sup>a</sup> | FVB, 9M<sup>a</sup> | HYB, 9M<sup>a</sup> |
| --- | --- | --- | --- | --- | --- | --- | --- | --- |
| Size | - | Weight [g] | 27<br>[25 28] | 31<br>[30 32] | 32<br>[30 35] | 30<br>[29 32] | 37<br>[34 38] | 45<br>[42 47] |
| **Sensorimotor behavior** |  |  |  |  |  |  |  |  |
| Visual behavior | Optomotor drum | Number of head turns | 8<br>[3 11] | 0<br>[0 0] | 7<br>[5 10] | 8<br>[5 11] | 0<br>[0 0] | 8<br>[5 9] |
| Motor endurance | Treadmill | Distance to failure [cm] | 228<br>[138 242] | 121<br>[73 197] | 222<br>[30 262] | 91<br>[49 120] | 41<br>[29 64] | 126<br>[59 164] |
| Balance | Rotarod | Latency to fall [s] | 260<br>[195 278] | 235<br>[194 246] | 140<br>[107 147] | 165<br>[141 186] | 123<br>[90 158] | 188<br>[139 221] |
| **Psychiatric-like behavior** |  |  |  |  |  |  |  |  |
| Anxiety-like behavior | Elevated plus maze | Contra-anxiety index | -0.59<br>[-0.66 -0.55] | -0.45<br>[-0.55 -0.39] | -0.34<br>[-0.55 0.22] | -0.56<br>[-0.75 -0.46] | -0.28<br>[-0.44 0.05] | -0.15<br>[-0.27 0.15] |
| Depressive-like behavior | Forced swim test | Freeze duration [s] | 124<br>[43 135] | 209<br>[95 257] | 201<br>[63 293] | 130<br>[86 172] | 0<br>[0 14] | 138<br>[85 189] |
| Exploration | Open field | Activity [%] | 0.23<br>[0.17 0.27] | 0.19<br>[0.15 0.22] | 0.28<br>[0.2 0.31] | 0.35<br>[0.33 0.38] | 0.3<br>[0.27 0.33] | 0.35<br>[0.28 0.37] |
| **Cognitive behavior** |  |  |  |  |  |  |  |  |
| Learning | Morris water maze | Latency to platform [s] | 21.7±4.3 s | 41.1±3.8 s | 19.3±3.9 s | 33.3±2.9 | 42.1±4 | 17.8±3 |
| Spatial long-term memory | MWM, probe day | Fraction of time in the target quadrant | 0.29<br>[0.23 0.36] | 0.21<br>[0.09 0.28] | 0.3<br>[0.19 0.39] | 0.3<br>[0.24 0.4] | 0.25<br>[0.15 0.31] | 0.55<br>[0.36 0.62] |

<sup>a</sup>Values indicate median [IQR] (inter-quartile interval) for every age group, strain, and metric. For learning on the MWM, values indicate mean±SEM.

**Table S3:** Mice used for electrophysiological recordings.

| Animal | Sex | Strain | Driver line<sup>a</sup> | Reporter line<sup>a</sup> | Viral vector | Age<sup>b</sup> [week] | Weight<sup>b</sup> [g] | Probe | Neocortex sessions | CA1 sessions |
| --- | --- | --- | --- | --- | --- | --- | --- | --- | --- | --- |
| mP23 | M | C57 | PV-Cre | Ai32 | - | 15 | 31.2 | Buzsaki32 | - | 14 |
| mDS1 | M | C57 | CaMKII-Cre | Ai32 | - | 14 | 25.7 | Dual-sided64 | - | 5 |
| mDS2 | F | C57 | CaMKII-Cre | Ai32 | - | 30 | 24.2 | Dual-sided64 | - | 21 |
| mC41 | M | HYB | CaMKII-Cre | Ai32 | - | 10 | 33.7 | Stark64 | 6 | 33 |
| mA234 | M | HYB | CaMKII-Cre | Ai32 | - | 16 | 30 | Buzsaki32 | 15 | 24 |
| mB142 | M | HYB | PV-Cre | - | DIO-ChR2 | 21 | 35.9 | Buzsaki32 | 1 | - |

<sup>a</sup>For C57 mice, the male parent was the reporter, and the female parent was the driver.
<sup>b</sup>At time of implantation.

**Table S4:**
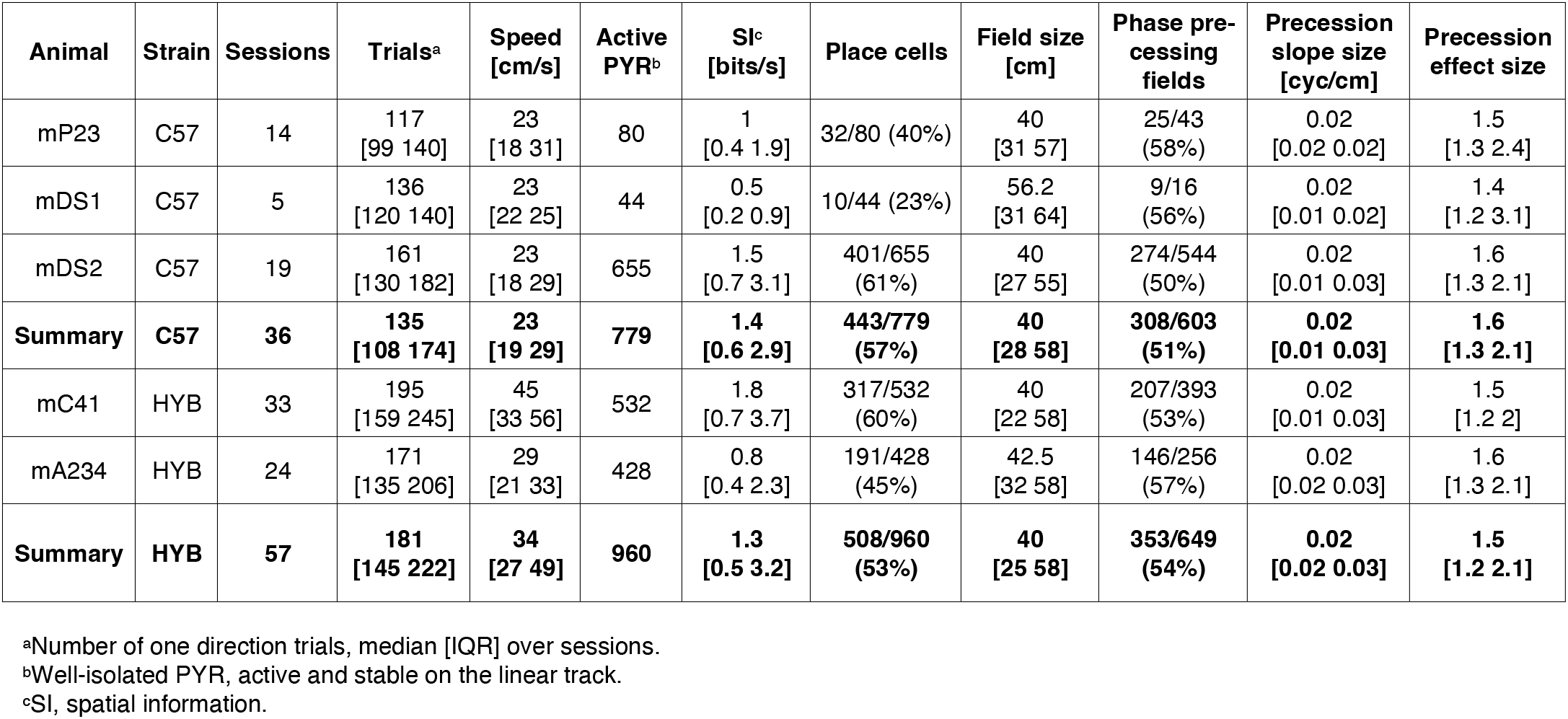
Linear track behavior and place coding in individual mice.

**Table S5:** Optogenetic activation of individual neurons in individual mice.

| Animal | Region | Sessions | Activated PYR<sup>a</sup> | Locally activated PYR<sup>b</sup> | Activated INT | Locally activated INT |
| --- | --- | --- | --- | --- | --- | --- |
| mA234 | CA1 | 24 | 493/1807 (27%) | 280/577 (48%) | 122/284 (43%) | 34/76 (45%) |
| mC41 | CA1 | 31 | 720/3761 (19%) | 442/950 (46%) | 119/782 (15%) | 58/197 (30%) |
| **Total** |  | **55** | **1213/5568 (22%)** | **722/1527 (47%)** | **241/1066 (23%)** | **92/273 (34%)** |
| mA234 | Neocortex | 15 | 653/1203 (54%) | 235/327 (72%) | 208/456 (46%) | 60/120 (50%) |
| mC41 | Neocortex | 6 | 82/343 (24%) | 39/82 (48%) | 21/61 (34%) | 6/11 (55%) |
| **Total** |  | **21** | **735/1564 (47%)** | **274/409 (67%)** | **229/517 (44%)** | **66/131 (50%)** |

<sup>a</sup>Well-isolated PYR with increased firing rate during optogenetic stimulation ( $p < 0.05$ , Poisson test), out of all PYR recorded in the same sessions.
<sup>b</sup>Light-activated PYR recorded on the illuminated shank.

## References

Abdeljalil, J., Hamid, M., Abdel-Mouttalib, O., Stéphane, R., Raymond, R., Johan, A., José, S., Pierre, C., Serge, P. (2005) The optomotor response: a robust first-line visual screening method for mice. Vision Res. 45, 1439–46.

Altman, P.L., Katz, D.D. (1979) Inbred and Genetically Defined Strains of Laboratory Animals, Part 1: Mouse and Rat. Bethesda, Federation of American Societies for Experimental Biology.

Beck, J.A., Lloyd, S., Hafezparast, M., Lennon-Pierce, M., Eppig, J.T., Festing, M.F., Fisher, E.M. (2000) Genealogies of mouse inbred strains. Nat. Genet. 24, 23–5.

Blednov, Y.A., Metten, P., Finn, D.A., Rhodes, J.S., Bergeson, S.E., Harris, R.A., Crabbe, J.C. (2005) Hybrid C57BL/6J x FVB/NJ mice drink more alcohol than do C57BL/6J-mice. Alcohol Clin. Exp. Res. 29, 1949–58.

Blednov, Y.A., Ozburn, A.R., Walker, D., Ahmed, S., Belknap, J.K., Harris, R.A. (2009) Hybrid mice as genetic models of high alcohol consumption. Behav. Genet. 40, 93–110.

Brekke, T.D., Steele, K.A., Mulley, J.F. (2018) Inbred or outbred? Genetic diversity in laboratory rodent colonies. G3 8, 679–86.

Bromley-Brits, K., Deng, Y., Song, W. (2011) Morris water maze test for learning and memory deficits in Alzheimer’s disease model mice. J. Vis. Exp. 53, 2920.

Brown, R.E., Wong, A.A. (2007) The influence of visual ability on learning and memory performance in 13 strains of mice. Learn. Mem. 14, 134–44.

Buhl, E., Whittington, M. (2007) Local circuits. In Andersen P, Morris R, Amaral D, Bliss T, O’Keefe J (Eds.), The hippocampus book. New York, Oxford University Press.

Can, A., Dao, D.T., Arad, M., Terrillion, C.E., Piantadosi, S.C., Gould, T.D. (2012) The mouse forced swim test. J. Vis. Exp. 59, 3638.

Castagné, V., Moser, P., Roux, S., Porsolt, R.D. (2011) Rodent models of depression: forced swim and tail suspension behavioral despair tests in rats and mice. Curr. Protoc. Pharmacol. 38, 5.8.1–11.

Christmas, A.J., Maxwell, D.R. (1970) A comparison of the effects of some benzodiazepines and other drugs on aggressive and exploratory behaviour in mice and rats. Neuropharmacology 9, 17–29.

Crawley, J.N. (1996) Unusual behavioral phenotypes of inbred mouse strains. Trends Neurosci. 19, 181–2.

Crawley, J.N. (2000) What’s wrong with my mouse? Behavioral phenotyping of transgenic and knockout mice. New York, Wiley-Liss.

Crawley, J.N. (2003) Behavioral phenotyping of rodents. Comp. Med. 53, 140–6.

Crawley, J.N. (2008) Behavioral phenotyping strategies for mutant mice. Neuron 57, 809–18.

Crowley, S.T., Kataoka, K., Itaka, K. (2018) Combined CatWalk Index: an improved method to measure mouse motor function using the automated gait analysis system. BMC Res. Notes 11, 263.

Fee, M.S., Mitra, P.P., Kleinfeld, D. (1996) Automatic sorting of multiple unit neuronal signals in the presence of anisotropic and non-Gaussian variability. J. Neurosci. Methods. 69, 175–88.

Fuchs, H. et al. (2011) Mouse phenotyping. Methods 53, 120–35.

Gaspar, N., Eichler, R., Stark, E. (2019) A novel low-noise movement tracking system with real-time analog output for closed-loop experiments. J. Neurosci. Methods, 318, 69–77

Grieco, F. et al. (2021) Measuring Behavior in the Home Cage: Study Design, Applications, Challenges, and Perspectives. Front. Behav. Neurosci. 15, 735387.

Jones, B.J., Roberts, D.J. (1968) The quantitative measurement of motor inco-ordination in naive mice using an accelerating rotarod. J. Pharm. Pharmacol. 20, 302–4.

Kadir, S.N., Goodman, D.F.M., Harris, K.D. (2014) High-dimensional cluster analysis with the masked EM algorithm. Neural Comput. 26, 2379–94.

Kafkafi, N., Golani, I., Jaljuli, I., Morgan, H., Sarig, T., Würbel, H., Yaacoby, S., Benjamini, Y. (2017) Addressing reproducibility in single-laboratory phenotyping experiments. Nat. Methods 14, 462–4.

Kempter, R., Leibold, C., Buzsáki, G., Diba, K., Schmidt, R. (2012) Quantifying circular-linear associations: hippocampal phase precession. J. Neurosci. Methods 207, 113–24.

Komada, M., Takao, K., Miyakawa, T. (2008) Elevated plus maze for mice. J. Vis. Exp. 22, 1088.

Mao, J.H., Langley, S.A., Huang, Y., Hang, M., Bouchard, K.E., Celniker, S.E., Brown, J.B., Jansson, J.K., Karpen, G.H., Snijders, A.M. (2015) Identification of genetic factors that modify motor performance and body weight using Collaborative Cross mice. Sci. Rep. 5, 16247.

Maroteaux, G. et al. (2012) High-throughput phenotyping of avoidance learning in mice discriminates different genotypes and identifies a novel gene. Genes Brain Behav. 11, 772–84.

Marques-Aleixo, I. et al. (2015) Physical exercise prior and during treatment reduces sub-chronic doxorubicin-induced mitochondrial toxicity and oxidative stress. Mitochondrion 20, 22–33.

Mekada, K., Abe, K., Murakami, A., Nakamura, S., Nakata, H., Moriwaki, K., Obata, Y., Yoshiki, A. (2009) Genetic differences among C57BL/6 substrains. Exp. Anim. 58, 141–9.

Mitchiner, J.C., Pinto, L.H., Vanable, J.W. Jr (1976) Visually evoked eye movements in the mouse (Mus musculus). Vision Res. 16, 1169–71.

Mogil, J.S. et al. (1999) Heritability of nociception I: responses of 11 inbred mouse strains on 12 measures of nociception. Pain 80, 67–82.

Moller, K.A., Berge, O.G., Hamers, F.P. (2000). Locomotor behaviour in normal and monoarthritic rats assessed by a new computer-assisted method. In Measuring Behavior Conference, Nijmegen, the Netherlands.

Morris, R.G. (1981) Spatial localization does not require the presence of local cues. Learn. Motiv. 12, 239–60.

Morris, R.G. (1984) Developments of a water-maze procedure for studying spatial learning in the rat. J. Neurosci. Methods 11, 47–60.

Moser, E.I., Kropff, E., Moser, M.B. (2008) Place cells, grid cells, and the brain’s spatial representation system. Annu. Rev. Neurosci. 31, 69–89.

Morse, H.C. III (1978) Origins of Inbred Mice. New York, Academic Press.

Namdar, I., Feldman, R., Glazer, S., Meningher, I., Shlobin, N.A., Rubovitch, V., Bikovski, L., Been, E., Pick, C.G. (2020) Motor effects of minimal traumatic brain injury in mice. J. Mol. Neurosci. 70, 365–77.

Nguyen, P.V., Abel, T., Kandel, E.R., Bourtchouladze, R. (2000) Strain-dependent differences in LTP and hippocampus-dependent memory in inbred mice. Learn. Mem. 7, 170–9.

Noked, O., Levi, A., Someck, S., Amber-Vitos, O., Stark, E. (2021) Bidirectional optogenetic control of inhibitory neurons in freely-moving mice. IEEE Trans. Biomed. Eng. 68, 416–27.

Noldus, L.P., Spink, A.J., Tegelenbosch, R.A. (2001) EthoVision: a versatile video tracking system for automation of behavioral experiments. Behav. Res. Methods Instrum. Comput. 33, 398–414.

O’Keefe, J., Dostrovsky, J. (1971) The hippocampus as a spatial map. Preliminary evidence from unit activity in the freely-moving rat. Brain. Res. 34, 171–5.

O’Keefe, J., Recce, M.L. (1993) Phase relationship between hippocampal place units and the EEG theta rhythm. Hippocampus 3, 317–30.

O’Leary, T.P., Gunn, R.K., Brown, R.E. (2013) What are we measuring when we test strain differences in anxiety in mice? Behav. Genet. 43, 34–50.

O’Leary, T.P., Savoie, V., Brown, R.E. (2011) Learning, memory and search strategies of inbred mouse strains with different visual abilities in the Barnes maze. Behav. Brain Res. 216, 531–42.

Olton, D.S., Becker, J.T., Handelmann, G.E. (1979) Hippocampus, space, and memory. Behav. Brain Sci. 2, 313–65.

Ouagazzal, A.M., Reiss, D., Romand, R. (2006) Effects of age-related hearing loss on startle reflex and prepulse inhibition in mice on pure and mixed C57BL and 129 genetic background. Behav. Brain Res. 172, 307–15.

Pachitariu, M., Steinmetz, N., Kadir, S, Carandini, M., Kenneth, D. H. (2016) Kilosort: realtime spike-sorting for extracellular electrophysiology with hundreds of channels. bioRxiv 061481. https://doi.org/10.1101/061481.

Pittler, S.J., Baehr, W. (1991) Identification of a nonsense mutation in the rod photoreceptor cGMP phosphodiesterase beta-subunit gene of the rd mouse. Proc. Natl. Acad. Sci. USA. 88, 8322–6.

Porsolt, R.D., Bertin, A., Jalfre, M.J. (1977) Behavioral despair in mice: a primary screening test for antidepressants. Arch. Int. Pharmacodyn. Ther. 229, 327–36.

Preisig, D.F., Kulic, L., Krüger, M., Wirth, F., McAfoose, J., Späni, C., Gan-tenbein, P., Derungs, R., Nitsch, R.M., Welt, T. (2016) High-speed video gait analysis reveals early and characteristic locomotor phenotypes in mouse models of neurodegenerative movement disorders. Behav. Brain Res. 311, 340–53.

Prevot, T.D., Misquitta, K.A., Fee, C., Newton, D.F., Chatterjee, D., Nikolova, Y.S., Sibille, E., Banasr, M. (2019) Residual avoidance: A new, consistent and repeatable readout of chronic stress-induced conflict anxiety reversible by antidepressant treatment. Neuropharmacology 153, 98–110.

Prut, L., Belzung, C. (2003) The open field as a paradigm to measure the effects of drugs on anxiety-like behaviors: a review. Eur. J. Pharmacol. 463, 3–33.

Rahn, R.M., Weichselbaum, C.T., Gutmann, D.H., Dougherty, J.D., Maloney, S.E. (2021) Shared developmental gait disruptions across two mouse models of neurodevelopmental disorders. J. Neurodev. Disord. 13, 10.

Rodgers, R.J., Dalvi, A. (1997) Anxiety, defence and the elevated plus-maze. Neurosci. Biobehav. Rev. 21, 801–10.

Rossant, C. et al. (2016) Spike sorting for large, dense electrode arrays. Nat. Neurosci. 19, 634–41.

Schmidt, R., Diba, K., Leibold, C., Schmitz, D., Buzsáki, G., Kempter, R. (2009) Single-trial phase precession in the hippocampus. J. Neurosci. 29, 1323241.

Schmitzer-Torbert, N., Jackson, J., Henze, D., Harris, K., Redish, A.D. (2005) Quantitative measures of cluster quality for use in extracellular recordings. Neuroscience 131, 1–11.

Shiotsuki, H., Yoshimi, K., Shimo, Y., Funayama, M., Takamatsu, Y., Ikeda, K., Takahashi, R., Kitazawa, S., Hattori, N. (2010) A rotarod test for evaluation of motor skill learning. J. Neurosci. Methods. 189, 180–5.

Silver, L.M. (1995) Mouse genetics: Concepts and appiications. New York, Oxford University Press.

Silva, A.J. et al. (1997) Mutant mice and neuroscience: recommendations concerning genetic background. Banbury Conference on genetic background in mice. Neuron 19, 755–59.

Simon, M.M. et al. (2013) A comparative phenotypic and genomic analysis of C57BL/6J and C57BL/6N mouse strains. Genome Biology. 14, 1–22.

Sittig, L.J., Carbonetto, P., Engel, K.A., Krauss, K.S., Barrios-Camacho, C.M., Palmer, A.A. (2016) Genetic background limits generalizability of genotype-phenotype relationships. Neuron 91, 1253–9.

Sloin, H.E., Levi, A., Someck, S., Spivak, L., Stark, E. (2022) High fidelity theta phase rolling of CA1 neurons. J. Neurosci. 42, 3184–96.

Stark, E., Koos, T., Buzsáki, G. (2012) Diode-probes for spatiotemporal optical control of multiple neurons in freely-moving animals. J. Neurophysiol. 108, 349–63.

Stark, E., Eichler, R., Roux, L., Fujisawa, S., Rotstein, H.G., Buzsáki, G. (2013) Inhibition-induced theta resonance in cortical circuits. Neuron 80, 126376.

Taketo, M. et al. (1991) FVB/N: an inbred mouse strain preferable for transgenic analyses. Proc. Natl. Acad. Sci. USA 88, 2065–9.

Terry, A.V. Jr (2009) Spatial navigation (water maze) tasks. In Buccafusco JJ (Ed.), Methods of Behavior Analysis in Neuroscience. Boca Raton, CRC Press.

Thomson, A.M., Radpour, S. (1991) Excitatory connections between CA1 pyramidal cells revealed by spike triggered averaging in slices of rat hippocampus are partially NMDA receptor mediated. Eur. J. Neurosci. 3, 587601.

Upchurch, M., Wehner, J.M. (1988) Differences between inbred strains of mice in Morris water maze performance. Behav. Genet. 18, 55–68.

de Visser, L., van den Bos, R., Kuurman, W.W., Kas, M.J., Spruijt, B.M. (2006) Novel approach to the behavioural characterization of inbred mice: automated home cage observations. Genes Brain Behav. 5, 458–66.

Võikar, V., Kõks, S., Vasar, E., Rauvala, H. (2001) Strain and gender differences in the behavior of mouse lines commonly used in transgenic studies. Physiol. Behav. 72, 271–81.

Vorhees, C.V., Williams, M.T. (2006) Morris water maze: procedures for assessing spatial and related forms of learning and memory. Nat. Protoc. 1, 848–58.

Wahlsten, D. et al. (2003) Different data from different labs: lessons from studies of gene-environment interaction. J. Neurobiol. 54, 283–311.

Waterston, R.H. et al. (2002) Initial sequencing and comparative analysis of the mouse genome. Nature 420, 520–62.

Zurita, E. et al. (2011) Genetic polymorphisms among C57BL/6 mouse inbred strains. Transgenic Res. 20, 481–89.

